# Vitamin D mitigates Inflammatory Bone Loss in Postmenopausal Osteoporosis via modulating the Gut-Immune-Bone axis

**DOI:** 10.64898/2026.06.29.735433

**Authors:** Asha Bhardwaj, Leena Sapra, Tamanna Sharma, Swati Rajput, Anurag Singh, Sumedha Yadav, Chaman Saini, Pradyumna K. Mishra, Bhavuk Garg, Vikrant Manhas, Pragya Shukla, Adarsh Wamanrao Barwad, Rupesh K. Srivastava

**Author notes:** Correspondence &.

## Abstract

Osteoporosis is a prevalent skeletal disorder characterized by deterioration of bone microarchitecture and loss of bone mineral density, leading to increased fracture risk and substantial health and economic burdens, particularly among older adults. Bone remodeling is orchestrated by a complex interplay of systemic and local regulators, among which vitamin D plays a central role in maintaining skeletal homeostasis. Although numerous studies have examined the effects of vitamin D on bone metabolism, outcomes have been inconsistent across populations, dosing regimens, and experimental models. To clarify the net skeletal impact of vitamin D, we investigated its effects in postmenopausal osteoporosis (PMO). Vitamin D (1,25-dihydroxyvitamin D3– active form of vitamin D) supplementation effectively prevented bone loss in ovariectomized mice, at both lower and higher concentrations. Mechanistically, vitamin D promoted osteoclast differentiation *in vitro*, consistent with its RANKL-dependent pro-osteoclastogenic activity, yet paradoxically conferred bone protection *in vivo*. This discrepancy was explained by vitamin D’s profound immunomodulatory effects, which reshaped both innate and adaptive immune responses to suppress osteoclast formation and function. Concurrently, vitamin D improved intestinal barrier integrity and restored gut microbial composition, thereby stabilizing the gut-immune-bone axis and reducing pro-resorptive inflammatory signaling. Together, these findings demonstrate that vitamin D prevents bone loss through the coordinated regulation of immune and gut homeostasis, reconciling its apparent pro-resorptive effects *in vitro* with its overall anti-resorptive outcomes *in vivo*. This integrated mechanism highlights immune-gut microbial modulation as a key mediator of vitamin D-induced bone preservation and supports the development of vitamin D as an immunotherapeutic adjunct for the prevention and management of PMO. Altogether, our findings for the first time dissect the paradox surrounding the osteoprotective property of vitamin D supplementation.

**Graphical Abstract:**
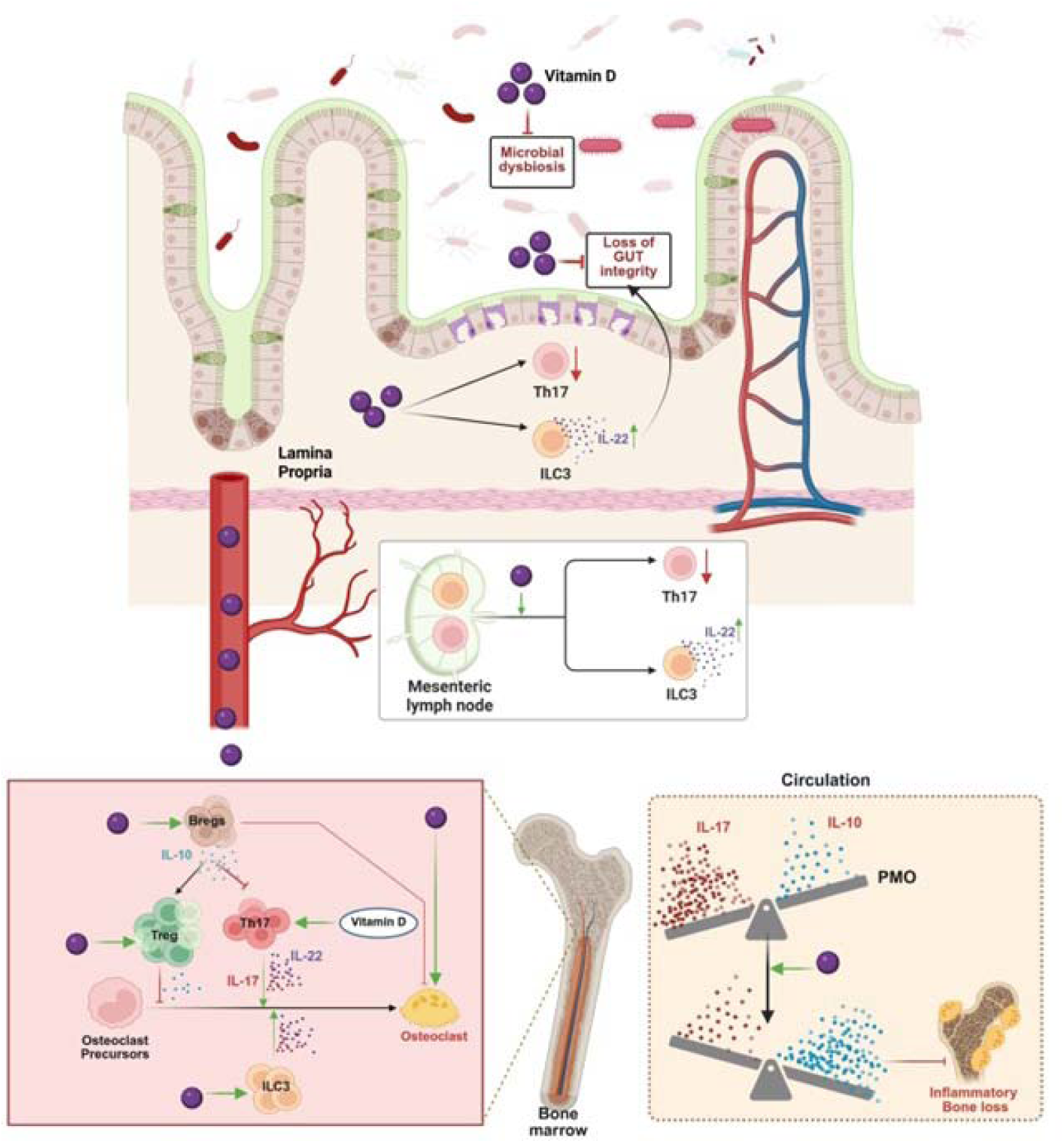
Vitamin D strengthens intestinal barrier integrity and restores gut microbial composition, which stabilizes the gut-bone axis and suppresses pro-resorptive inflammatory signaling. Vitamin D enhances osteoclastogenesis both directly and through ILC3– and Th17-mediated pathways, while inhibiting osteoclastogenesis via Treg– and Breg-dependent mechanisms. In circulation, vitamin D maintains the homeostasis between IL-10 and IL-17 cytokines. Collectively, these results indicate that vitamin D helps prevent bone loss by orchestrating the immune system and maintaining gut homeostasis.

## 1.0 Introduction

Osteoporosis is one of the most prevalent bone disorders worldwide, characterized by low bone mass and deterioration of bone microarchitecture, leading to decreased bone strength and an increased risk of fractures. It affects more than 500 million people globally, with over one in two women and one in four men above the age of 50 experiencing an osteoporotic fracture (International Osteoporosis Foundation, 2025). This imposes a substantial health and economic burden on society. Although several therapeutic agents are currently available for the prevention and treatment of osteoporosis, their long-term use is often limited by side effects and suboptimal efficacy, highlighting the urgent need for safer and more effective interventions.

Osteoporosis develops from an imbalance in bone remodeling, where osteoclast-mediated bone resorption exceeds osteoblast-driven bone formation. This dynamic process is tightly regulated by a complex interplay of biochemical and mechanical factors, including parathyroid hormone (PTH), estrogen, glucocorticoids, insulin-like growth factor (IGF), bone morphogenetic proteins (BMPs), and transforming growth factor-β (TGF-β)^1^. Among these, vitamin D is a key regulator of skeletal homeostasis. It facilitates intestinal calcium and phosphate absorption by stimulating calcium channel activity and promoting the synthesis of calcium-binding proteins in enterocytes. Consequently, vitamin D deficiency impairs mineralization, leading to bone loss and skeletal abnormalities such as osteoporosis^2^.

Vitamin D exerts both direct and indirect effects on bone cells, influencing the activity of osteoblasts and osteoclasts, and is known to reduce the risk of fractures. Vitamin D deficiency is highly prevalent among postmenopausal osteoporosis (PMO) patients^3^. Regular consumption of milk enriched with calcium and vitamin D has been shown to improve bone mineral density (BMD), particularly in the femoral neck of postmenopausal women. Meta-analyses further demonstrate that combined supplementation of vitamin D and calcium effectively reduces fracture incidence, suggesting that the bone-protective effects of vitamin D are most pronounced when administered together with calcium^4^. Animal studies provide important mechanistic insights into the role of vitamin D in skeletal health. In mice with thyrotoxicosis, vitamin D treatment suppresses bone resorption and enhances BMD and trabecular bone microarchitecture by increasing the osteoprotegerin (OPG) to receptor activator of nuclear factor kappa-Β ligand (RANKL) ratio and activating the Wnt/β-catenin signaling pathway^5^. Similarly, in rat models exposed to microgravity, vitamin D in combination with calcium prevents bone mineral loss^6^. In rabbit models of inflammation-induced osteoporosis, high doses of vitamin D suppress bone resorption and inflammation, but significant improvement in mineralization occurs only when calcium supplementation is provided concurrently^7^. Several studies also indicate that vitamin D’s effects are dose-dependent. Higher doses of vitamin D have been associated with approximately 20% or greater reductions in nonvertebral fracture risk among individuals aged 65 years and older^8,9^. In animal studies, supplementation with 20,000 IU/kg vitamin D□ during early adulthood led to the development of stronger, more ductile bones compared to standard doses^10^. However, not all findings are consistent. Long-term calcium and vitamin D supplementation in ovariectomized rat models failed to produce significant improvements in BMD or bone microarchitecture, despite preventing weight gain^11^. Similarly, high-dose vitamin D therapy modestly enhanced calcium absorption but did not consistently improve BMD, muscle strength, or fall reduction in PMO women^12^. Mechanistically, the active metabolite 1,25-dihydroxyvitamin D□ can induce RANKL expression, promote osteoclast differentiation, and potentially enhance bone resorption^13^. Excessive vitamin D intake may also lead to adverse outcomes, particularly in females, underscoring the importance of precise dose optimization. Furthermore, there is insufficient evidence to support vitamin D alone (without calcium) as an effective means of preventing bone loss^4^. Overall, evidence supporting vitamin D as a standalone therapy for preventing bone loss remains insufficient. The dual effects of vitamin D, both promoting osteoclastogenesis and inhibiting bone loss through regulatory mechanisms, reflect its complex and context-dependent influence on bone remodeling. Therefore, despite substantial research, the independent role of vitamin D (without calcium) in bone maintenance remains unclear. Moreover, vitamin D exhibits immunomodulatory effects that influence bone health, an aspect often overlooked in analyses of its skeletal impacts^14–18^. Further research is therefore essential to clarify optimal dosing, timing, and underlying mechanisms to establish its therapeutic role in PMO.

Thus, in the present study, we investigated the effects of vitamin D supplementation on bone health using an ovariectomy (ovx)-induced osteoporosis pre-clinical mouse model. Mice were supplemented with two different concentrations of vitamin D to assess its dose-dependent effects on bone health, thereby examining the impact of both low and high doses of vitamin D. Our findings revealed that both doses of vitamin D showed bone-protective effects. Surprisingly, our findings further highlight the multifaceted nature of vitamin D: although it promotes osteoclastogenesis *in vitro*, of note, its dominant immunomodulatory role safeguards against bone loss. In parallel, vitamin D improved gut barrier integrity and mitigated dysbiosis, aligning with a gut-bone axis in which epithelial permeability and microbial composition modulate systemic osteoclastogenic cues. Taken together, these data indicate that vitamin D preserves bone in estrogen deficiency not merely by supporting mineral metabolism, but through coordinated immunomodulation and maintenance of the gut-immune-bone axis. This integrated mechanism reconciles the apparent paradox of pro-osteoclastogenic effects *in vitro* with net anti-resorptive outcomes *in vivo* and nominates immune–microbial pathways as leverage points for optimizing vitamin D-based strategies in osteoporosis prevention.

## 2.0 Materials & Methods

### 2.1 Animals

Female C57BL/6J mice (8-10 weeks old) were maintained under specific pathogen-free conditions at the Central Animal Facility, All India Institute of Medical Sciences (AIIMS), New Delhi, India. Animals were housed under a 12-hour light/dark cycle with ad libitum access to water and a standard chow diet containing adequate calcium sources, including dicalcium phosphate (1250 g/100 kg of diet) and calcium carbonate (555 g/100 kg of diet). The *in vivo* study comprised four groups, each containing six animals: sham-operated (control), ovariectomized (ovx), ovx treated with a low dose of 1,25-dihydroxyvitamin D□ (V1; 2□ng/mouse/day), and ovx treated with a higher dose (V2; 4□ng/mouse/day). The body weight of animals was recorded at regular intervals (Day 0, 15, 30, and 45) during the experimental period. After 45□days of treatment, animals were euthanized using carbon dioxide asphyxiation, and samples, including blood, bones, mesenteric lymph nodes (MLNs), and colon tissues, were collected for subsequent analyses. All procedures were performed following the guidelines for the care and use of laboratory animals and were approved by the Institutional Animal Ethics Committee (IAEC), AIIMS, New Delhi, India (approval number: 383/IAEC-1/2022).

### 2.2 Antibodies and reagents

For flow cytometry, the following antibodies and kits were procured from BD, Biolegend and eBioscience: CD3ε (553059), CD4 (553044), CD19 (553784), F4/80 (13-4801-82), NK1.1 (553163), FCεRI (13-5898-82), Gr-1 (553124), CD5 (553018), CD11c (553800), Ter119 (553672), streptavidin microbeads (557812), PE-Cy7 anti-mouse-CD3 (100220), APC anti-mouse-IL-22 (17-7222-80), BV786 anti-mouse-IL-17 (506928), PE anti-mouse-RORγT (12-6988-82), PerCp-Cy5.5 anti-mouse-CD19 (45-0193-82), BV421 anti-mouse-IL-10 (505022), APC anti-mouse-CD1d (17-0011-82), PECy7 anti-mouse-CD5 (25-0051-81), APC anti-mouse-Foxp3 (17-5773-82), BV711 anti-mouse-Tbet (644820), BV605 anti-mouse-interferon (IFN)-γ (505840), FITC anti-mouse-GATA 3 (53-9966-42), Foxp3/transcription factor staining buffer (0-5523-00), red blood cell (RBC) lysis buffer (00-4300-54) and monensin (420701). Phorbol myristate acetate (PMA) (P1585), ionomycin (10634), and acid phosphatase leukocyte (TRAP) kit (387A) were procured from Sigma-Aldrich (USA). Macrophage colony-stimulating factor (MCSF) (300-25) and RANKL (310-01) were procured from PeproTech (USA). α-Minimal essential medium (MEM) and Roswell Park Memorial Institute (RPMI)-1640 medium were obtained from Gibco (Thermo Fisher Scientific, USA). The following ELISA kits were brought from BD (USA): Mouse-tumour necrosis factor (TNF)-α, Mouse-interleukin (IL)-6, Mouse-IL-17, Mouse-IL-10, Mouse transforming growth factor (TGF)-β, and Mouse-IFNγ.

### 2.3 Purification and activation of innate lymphoid cell (ILC)-3

Purification and activation of ILC3 followed established protocols from our prior publication^19^. Briefly, single-cell suspensions were generated from spleens and subjected to RBC lysis before lineage depletion. The remaining cells were incubated for 30 minutes at 4°C with a biotinylated mouse ILC enrichment cocktail consisting of lineage markers (CD3, CD19, F4/80, NK1.1, FCεRI, Gr-1, CD5, CD11c, TER119). After staining, cells were washed with 1X PBS and further incubated with streptavidin-conjugated magnetic particles plus DM for 30 minutes at 4°C. The suspension was then placed in a magnetic separation unit, and the lineage-negative fraction enriched for ILCs was collected and evaluated for purity by flow cytometry, which consistently exceeded 90%. Enriched ILCs (50,000 cells per well) were seeded in 96-well plates and cultured for 24 hours at 37°C in a humidified atmosphere containing 5% CO□ in the presence or absence of ILC3-supporting cytokines (IL-2 and IL-7 at 10 ng/ml) and stimulatory cytokines (IL-23 and IL-1β at 50 ng/ml), with parallel conditions including or lacking vitamin D (50 nM). Following culture, the frequency and phenotype of ILC3s were determined by flow cytometric analysis.

### 2.4 Osteoclast differentiation and tartrate-resistant acid phosphatase (TRAP) staining

Osteoclast differentiation and TRAP staining followed established protocols from our prior publication^20^. Briefly, bone marrow cells (BMCs) were isolated from the femurs and tibiae of 8-12-week-old female C57BL/6 mice, followed by RBC lysis using 1X RBC lysis buffer. The cells were then plated in T25 flasks and cultured overnight in α-MEM containing 10% heat-inactivated fetal bovine serum (FBS) and M-CSF (35 ng/ml). The following day, non-adherent cells were collected and seeded at 50,000 cells per well in 96-well plates in osteoclastogenic medium supplemented with M-CSF (30 ng/ml) and RANKL (60 ng/ml), in the presence or absence of vitamin D (50 nM), IL-22 (10, 20, or 50 ng/ml), or vitamin D combined with short-chain fatty acids (SCFAs; 0.5 mM each of acetate, propionate, butyrate, and valerate). On day 3, half of the culture medium was replaced with fresh complete α-MEM containing the respective supplements. To evaluate osteoclast differentiation, cells were subjected to TRAP staining at the end of the culture period. After two washes with 1X PBS, cells were fixed with a citrate-acetone-3.7% formaldehyde solution for 10 minutes at 37°C and then incubated with TRAP staining solutions A and B according to the manufacturer’s instructions at 37°C in the dark for 10-15 minutes. Multinucleated TRAP-positive cells containing three or more nuclei were defined as mature osteoclasts and were counted and imaged using an inverted microscope (ECLIPSE TS100, Nikon). The TRAP-positive cell area was quantified using ImageJ software (NIH, USA).

### 2.5 Co-culture of ILC3 with BMCs for Osteoclastogenesis

Co-culture of ILC3 with BMCs for osteoclastogenesis followed established protocols from our prior publication^19^. Briefly, BMCs were isolated from femurs and tibiae of 8-12-week-old female C57BL/6 mice, followed by RBC lysis with 1X RBC lysis buffer. Cells were cultured overnight in T25 flasks using α-MEM supplemented with 10% heat-inactivated FBS and M-CSF (35 ng/ml). Non-adherent cells were then harvested and plated at 50,000 cells per well in 96-well plates with osteoclastogenic medium containing M-CSF (30 ng/ml) and RANKL (60 ng/ml), together with unprimed or vitamin D-primed ILC3s (BMCs: ILC3 at 1:1 or 1:5) for 4 days. Half the medium was replaced with fresh complete medium containing supplements on day 3. Osteoclast formation was assessed by TRAP staining at culture endpoint. For the IL-22 neutralization experimental setup, neutralizing monoclonal antibody against IL-22 (10 μg/ml) was added in the cocultures of BMCs and vitamin D-primed ILC3s at a 1:1 ratio in the presence of M-CSF and RANKL. After 4 days of incubation, cells were processed for evaluating osteoclastogenesis via TRAP staining.

### 2.6 Tregs and Th17 cell differentiation and coculture with BMCs for osteoclastogenesis

Tregs and Th17 cell differentiation followed established protocols from our prior publication^21^. Briefly, naïve CD4^+^ T cells were isolated by negative selection using a T cell enrichment kit (BD Biosciences) from lymph nodes of 8-12-week-old mice and plated on a culture plate coated with anti-CD3 (10 µg/ml) and anti-CD28 (2 µg/ml). For Treg differentiation, cells received TGF-β1 (5 ng/ml), IL-2 (10 ng/ml), anti-IL-4 (5 µg/ml), and anti-IFNγ (5 µg/ml), with or without vitamin D (50 nM), over 5 days. Th17 differentiation used anti-IL-4 (10 µg/ml), anti-IFNγ (10 µg/ml), TGF-β1 (2 ng/ml), IL-6 (30 ng/ml), and IL-23 (20 ng/ml), again with or without vitamin D (50 nM), for 5 days. On day 5, cells were collected for flow cytometry to quantify Treg and Th17 populations. To test their impact on osteoclastogenesis, BMCs (prepared as described previously) were cocultured with unprimed or vitamin D-primed Tregs/Th17 cells at ratios of 1:1 and 1:5 (BMCs: Treg/Th17) in the presence of M-CSF and RANKL. Multinucleated osteoclasts were then identified by TRAP staining.

### 2.7 Breg differentiation and coculture with BMCs for osteoclastogenesis

Breg differentiation and coculture with BMCs for osteoclastogenesis followed established protocols from our prior publication^20^. Briefly, splenic B cells were purified from C57BL/6 mice by magnetic negative selection following the manufacturer’s protocol (BD Biosciences): post-RBC lysis, single-cell suspensions were incubated with a biotinylated B cell enrichment cocktail for 20-30 min at 4°C, washed, treated with streptavidin particles plus DM for 30 min at 4°C, and the lineage-negative fraction collected after magnet separation with >95% purity confirmed by flow cytometry. Purified B cells (2 × 10□/well in 1 ml) were then cultured for 24 h at 37°C in 5% CO□ in 24-well plates with or without vitamin D (50 nM). Unprimed and vitamin D-primed Bregs were washed 3 times with PBS before coculture with BMCs (prepared as described previously) at ratios of 1:5 and 1:10 (BMCs: Bregs) in M-CSF/RANKL-containing medium, followed by TRAP staining to quantify multinucleated osteoclasts.

### 2.8 Flow Cytometry

Flow cytometry followed established protocols from our prior publication^22^. Briefly, cells from bone marrow, MLN and intestine were harvested, processed into single-cell suspensions in RPMI-1640 medium, and plated at 1 × 10^6^ cells per well in 96-well plates. Stimulation was performed with PMA (50 ng/ml) and ionomycin (1 µg/ml) for 5 hours, with monensin added during the final 2.5 hours to block protein transport. Cells were then surface-stained with fluorochrome-conjugated antibodies against CD3 (for ILCs, Th1, Th2, Th17), CD19/CD1d/CD5 (Bregs), and CD4 (Tregs) for 30-45 minutes on ice in the dark with gentle tapping every 5 minutes. Following washing, cells were fixed/permeabilized using 1X eBioscience™ Foxp3/transcription factor staining kit and stained intracellularly with antibodies against FoxP3, RORγt, IL-10, IL-17, T-bet, GATA-3, IFN-γ, and IL-22. Samples were acquired on a BD FACS Symphony flow cytometer (BD Biosciences) and analyzed with FlowJo v10.8.1 software (BD Biosciences). Gating strategies for ILCs, T cell subsets, and Bregs are detailed in **Supplementary Figures 1-3.**

### 2.9 Postmenopausal osteoporotic preclinical mice model (PMO)

Bilateral ovariectomy was performed in the ovx group under intraperitoneal anaesthesia. A 1 cm ventral incision exposed the peritoneal cavity, allowing visualization of both ovaries via the uterine horns. Fallopian tubes were ligated with sterile sutures proximal to the ovaries, which were then excised using surgical scissors. The peritoneum was closed with continuous sutures, followed by skin closure; mice received postoperative care for 7 days.

### 2.10 Cytokine analysis by Enzyme-Linked Immunosorbent Assay (ELISA)

Blood was collected by retro-orbital bleeding, allowed to clot for 45 minutes at room temperature, and then centrifuged at 4000 rpm for 20 minutes to obtain serum, which was stored at –80°C until analysis. Serum levels of osteoclastogenic cytokines (TNF-α, IL-6, IL-10, IL-17, TGF-β, and IFN-γ) were quantified by ELISA.

### 2.11 Micro-computed Tomography (µCT)

Bone samples were fixed in 4% paraformaldehyde for 3-4 days and then transferred to 1X PBS for storage. Micro-CT analysis of lumbar vertebra (LV)-5, femur, and tibia was conducted using a SkyScan 1176 scanner at 100 kV and 100 µA with a 0.5 mm aluminium filter (exposure time: 590 ms), acquiring 1800 projections at 6.93 µm resolution. Projection data were reconstructed with NRecon software, followed by morphometric analysis of trabecular and cortical bone parameters using CT Analyzer v1.02 (SkyScan); 2D/3D rendering of cortical bone was performed with Batman software.

### 2.12 Hematoxylin, Eosin (H&E) and TRAP staining

Femurs from mice were dissected, cleaned of surrounding soft tissues, and fixed in 4% paraformaldehyde for 48-72 h. Samples were decalcified in 10% EDTA (pH 7.4) for 2 weeks, followed by graded dehydration and paraffin embedding. Serial sections (4-5 μm) were prepared using a microtome. For histological evaluation, sections were stained with (H&E) to assess bone microarchitecture, including trabecular structure and marrow region. For osteoclast detection, TRAP staining was performed. TRAP-positive multinucleated cells were identified as osteoclasts. Images were captured using a light microscope, and quantitative analysis was performed using ImageJ software. For histological evaluation of non-skeletal tissues, liver, lung, kidney, and heart samples were fixed in 10% formaldehyde at room temperature overnight, followed by paraffin embedding, sectioning, and H&E staining as described above.

### 2.13 16S rRNA microbial community analysis

16S rRNA microbial community analysis followed established protocols from our prior publication^22^. Fecal pellets collected from sham and ovx mice on day 45 were immediately snap-frozen in sterile tubes at –80°C for metagenomic analysis. Metagenomic DNA was extracted using the QIAamp DNA Stool Mini Kit (QIAGEN) per manufacturer instructions, with DNA quality verified by NanoDrop A260/280 ratios. Qualified samples underwent 16S rRNA amplicon generation and Nextera XT library preparation (Illumina) for sequencing on the MiSeq platform. The QIIME2 pipeline analyzed microbiota composition: raw reads were quality-filtered with Trimmomatic v0.38 (sliding window 20 bp, min length 100 bp) to remove adapters, >5% ambiguous bases, and low-quality sequences (>10% bases <Q25), then merged paired-end reads with FLASH and denoised/chimera-filtered using DADA2. Amplicon sequence variants received SILVA-based taxonomic assignment via q2-feature-classify, with alpha diversity (observed ASVs) and statistical comparisons computed in QIIME2.

### 2.14 Gut permeability assay (Evans Blue Assay)

Mice received tail vein injections of Evans Blue dye (30 mg/kg body weight). After 30 minutes, animals were euthanized, large intestines were excised and incubated in formamide at 72°C for 24 hours, and dye extravasation was quantified by absorbance of the supernatant at 620 nm.

### 2.15 Study Subjects

Sixty-eight individuals were recruited for the study. Bone mineral density was assessed using dual-energy X-ray absorptiometry (DEXA), and participants were categorized as healthy (T-score > −1.0), osteopenic (T-score –1.0 to –2.5), or osteoporotic (T-score < –2.5). Individuals with a history of smoking, alcohol use, diabetes, endocrine disorders, or hypertension were excluded. Participants receiving anti-osteoporotic therapy, glucocorticoids, hormone replacement therapy, oral contraceptives, or other medications known to influence bone metabolism were also excluded, as were pregnant or lactating women. The study was conducted following approval from the Institute Ethics Committee for Postgraduate Research at the All India Institute of Medical Sciences, New Delhi (IECPG-482). Serum samples from all participants were analysed for vitamin D levels and for IL-10, IL-17, and C-terminal telopeptide-1 (CTX-1) using ELISA. In a subset of ten participants, peripheral blood mononuclear cells were isolated for flow cytometric evaluation of Tregs, Bregs and Th17 cells.

### 2.16 qPCR

Gene expression analysis was performed by RT-PCR using a QuantStudio 5 system (Applied Biosystems). cDNA samples from each experimental group were run in duplicate with custom primers targeting claudin-1, occludin, IL-17A, IL-22, VDR, 24-hydroxylase, and 1-α-hydroxylase, normalized to GAPDH as the reference gene. Reactions contained 25 ng cDNA, 2X SYBR Green PCR Master Mix (Promega), and primers in each well; relative expression was calculated from normalized threshold cycle (Ct) values.

### 2.17 Statistical Analysis

Comparisons between experimental groups were performed using paired or unpaired Student’s t-tests or one-way ANOVA, followed by Dunnett’s test, as appropriate for the study design. Results are presented as mean values with the corresponding standard error of the mean (SEM). A p-value of 0.05 or less was considered statistically significant, with significance levels denoted as *p < 0.05, **p < 0.01, ***p < 0.001, and ****p < 0.0001 for the indicated comparisons.

## 3.0 Results

### 3.1 Vitamin D ameliorates bone loss in PMO

First, we investigated the association between serum vitamin D levels and PMO. To this end, PMO women were recruited, and BMD was assessed using DEXA scan. Serum samples from these subjects were analysed for vitamin D and CTX-1 levels. Interestingly, vitamin D levels showed a positive correlation with BMD but a negative correlation with both T-score and CTX-1 levels, indicating a strong association between vitamin D deficiency and osteoporosis **(Figure 1A-C).** To further elucidate the mechanistic role of vitamin D in bone health and to assess whether vitamin D supplementation could improve bone integrity, we next evaluated its direct effect using an *in vivo* mouse model. To accomplish the same, mice were divided into four groups: sham (control), ovx (ovariectomized), and ovx + V1 (2 ng/mice/day) and ovx + V2 (4 ng/mice/day). On day 45, mice were sacrificed, and tissues were collected for various analyses **(Figure 1D).** To evaluate the safety profile of vitamin D supplementation, potential adverse effects were assessed in both treatment groups. No significant differences in body weight were observed among control, ovx, and vitamin D-treated groups throughout the study period **(Supplementary Figure 4A).** In addition, histological examination of major non-skeletal organs, including liver, kidney, and lung, revealed no detectable pathological alterations **(Supplementary Figure 4B).** These findings indicate that vitamin D administration at the tested doses was well tolerated and did not induce observable systemic toxicity. Next, to assess the effect of vitamin D supplementation on bone health, μ-CT analysis was performed. μ-CT results demonstrated a significantly more deteriorated microarchitecture of the lumbar vertebra (LV)-5, femur, and tibia in the ovx group compared with the sham group. Notably, vitamin□D administration (both V1 and V2) markedly improved the 3D microarchitecture of these bones in ovx mice **(Figure□1E).** Histomorphometric analysis of the LV-5 vertebra revealed a marked loss of bone volume and trabecular deterioration in the ovx group compared to the sham group **(Figure□2A).** Interestingly, vitamin□D supplementation significantly increased trabecular thickness (Tb.Th,□p□<□0.05) and decreased trabecular separation (Tb.Sp,□p□<□0.05) compared with the ovx group **(Figure□2A).** Examination of histomorphometric parameters of femoral and tibial bones showed that V1 and V2 treatment significantly improved bone volume by tissue volume (BV/TV, p□<□0.05) and trabecular number (Tb.N, p□<□0.05) of the tibial trabecular bone. V2 administration also further significantly improves the periosteal area (T.Ar, p□<□0.05) and periosteal perimeter (T.Pm, p□<□0.05) of the femoral cortical bone **(Supplementary□Figure□5).** Moreover, the BMD of the femoral trabecular bone was significantly reduced in ovx mice compared with the sham group, whereas V2 treatment effectively restored it to near-normal levels **(Figure 2B).** To further evaluate bone quality, we assessed the bending strength of femurs. Interestingly, the energy to failure was markedly decreased in ovx mice relative to sham controls; however, this mechanical parameter was significantly restored in the ovx + V1 and ovx + V2 groups **(Figure 2B).**

**Fig. 1:**
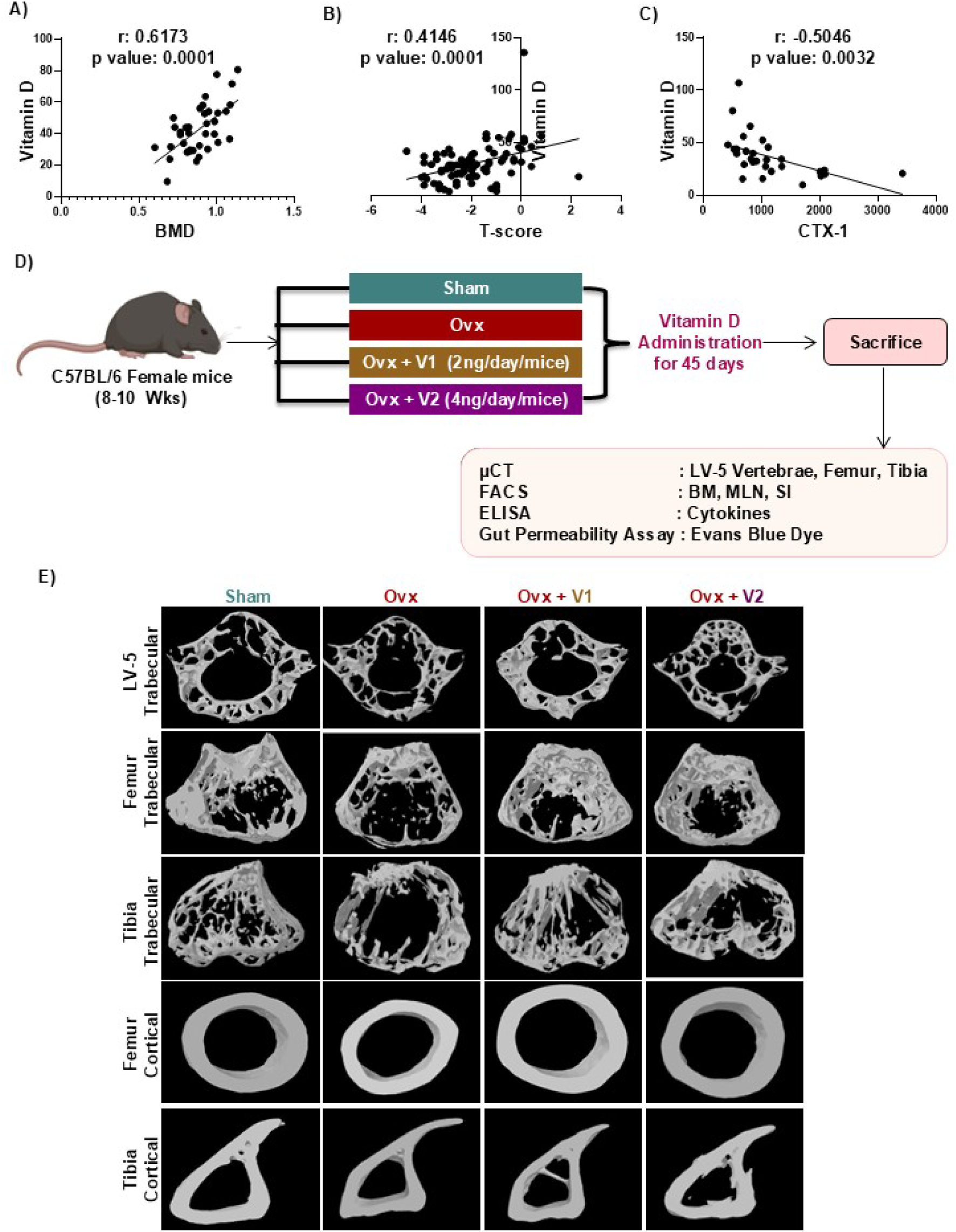
Experiment work plan for *in vivo* experiment. **(A)** Correlation between the bone mineral density (BMD) and the vitamin D levels in the human subjects. (**B)** Correlation between the T score (lumbar spine) and the vitamin D levels in the human subjects. **C)** Correlation between the C-terminal telopeptide (CTX) levels and the vitamin D levels in the human subjects. (**D)** Mice were divided into 4 groups: sham, ovx, ovx + V1 and ovx + V2. After 45 days, the mice were sacrificed and analysed for various parameters. (**E)** 3-D µ-CT reconstruction images of the LV-5 trabecular, femur trabecular, tibia trabecular, femur cortical, and tibia cortical regions of the sham, ovx, ovx + V1 and ovx + V2. groups.

**Fig. 2:**
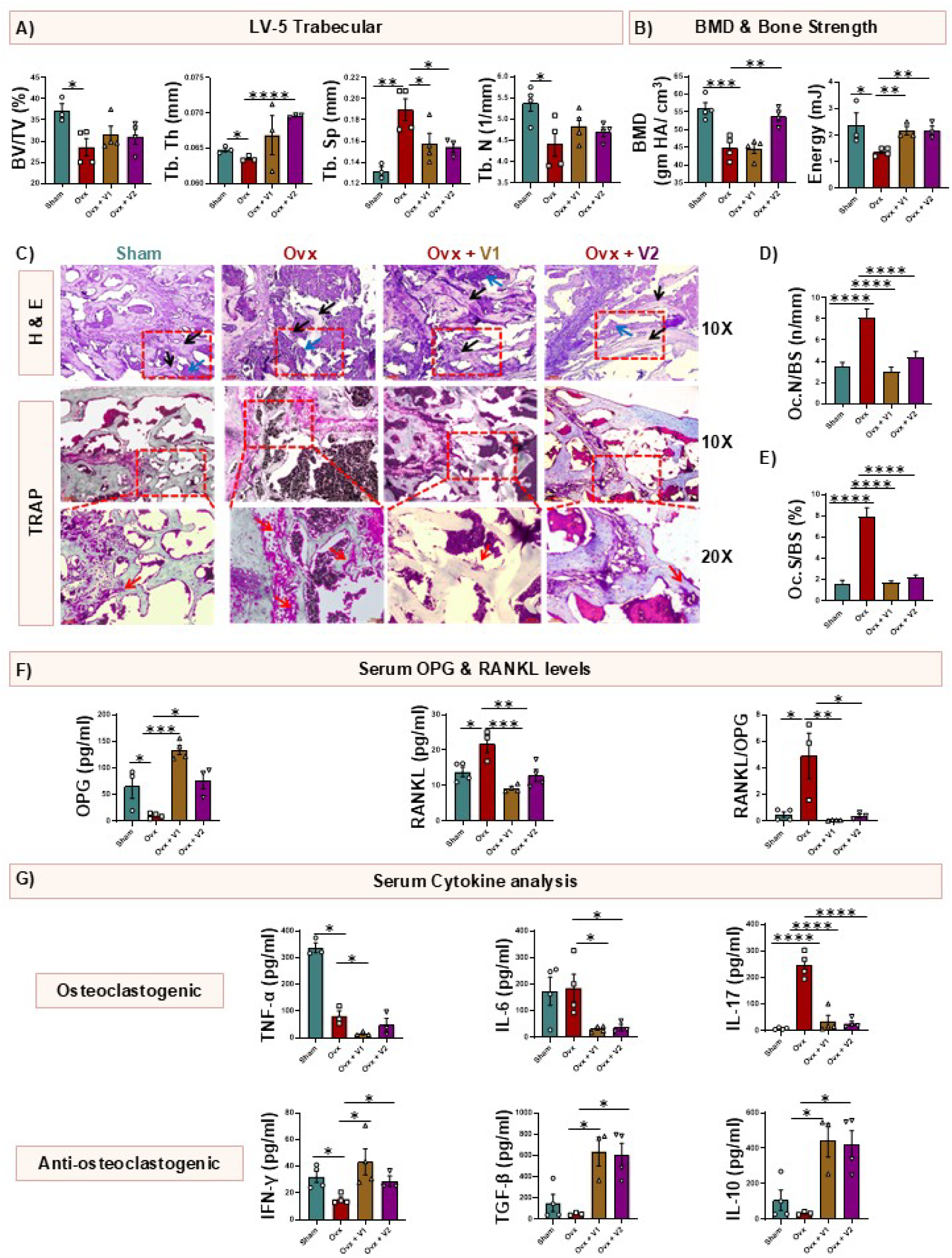
Vitamin D administration prevents bone loss in ovx mice. **(A**) Histomorphometric parameters of the LV-5 trabecular bone. (**B)** Graphical representation of bone mineral density (BMD) of the femur trabecular region and three-point bending test of femur diaphysis representing energy (mJ). (**C)** Hematoxylin and eosin (H&E) and tartrate-resistant acid phosphatase (TRAP) staining were performed on bone sections from all experimental groups (n=3). The top panel shows H&E-stained sections of the femoral metaphysis, highlighting trabecular bone (black arrows) and marrow regions (blue arrows); images were captured at 10X magnification. The middle and bottom panels display TRAP-stained sections of the femoral metaphysis, where pink/purple-stained cells represent osteoclasts, with red arrows indicating multinucleated osteoclasts; images were acquired at 10X and 20X magnifications. Quantitative analysis of **(D)** osteoclast number per bone surface (Oc.N/BS) and **(E)** osteoclast surface per bone surface (Oc.S/BS, %) was performed from TRAP-stained sections at 10X magnification using ImageJ software. **(F)** Serum osteoprotegerin (OPG) and receptor activator of nuclear factor κB ligand (RANKL) levels. (**G)** Osteoclastogenic and anti-osteoclastogenic cytokines were analyzed in serum samples of mice by ELISA. BV/TV, bone volume/tissue volume ratio; Tb.Th.– trabecular thickness; Tb.Sp.– trabecular separation; Tb.N– trabecular number; TNF– tumour necrosis factor; IL– interleukin; TGF– transforming growth factor. Data are represented as Mean ± SEM. The results were evaluated by one-way ANOVA followed by Dunnett’s test. Statistical significance was defined as *p < 0.05, **p < 0.01, ***p < 0.001 and ****p < 0.0001, concerning the indicated mouse group.

To further evaluate the protective effect of vitamin D against ovx-induced bone loss, H&E and TRAP staining were performed on bone sections from all groups. H&E staining revealed marked deterioration of trabecular architecture in ovx mice. In contrast, vitamin D-treated groups exhibited preservation of trabecular structure **(Figure 2C).** Consistently, TRAP staining showed a significant increase in TRAP-positive multinucleated osteoclasts along trabecular surfaces in ovx mice **(Figure 2C),** reflecting enhanced osteoclastogenesis, which was further supported by elevated osteoclast number per bone surface (Oc.N/BS) and osteoclast surface per bone surface (Oc.S/BS) **(Figure 2D-E).** Vitamin D treatment markedly reduced TRAP-positive cells along with a corresponding decrease in Oc.N/BS and Oc.S/BS, indicating suppression of osteoclast activity. **(Figure 2C-E).** Collectively, these findings demonstrate that vitamin D preserves bone integrity by inhibiting osteoclast-mediated bone resorption and maintaining trabecular architecture. At the molecular level, osteoprotegerin (OPG) levels were significantly decreased in ovx mice, while both V1 and V2 treatments markedly elevated OPG expression **(Figure 2F).** Conversely, RANKL levels were substantially increased in ovx mice but normalized following V1 and V2 administration **(Figure 2F).** RANKL to OPG ratio was significantly enhanced in the ovx mice but markedly reduced on the V1 and V2 administration **(Figure 2F).** Consistent with these findings, serum cytokine profiling revealed elevated levels of osteoclastogenic cytokines (TNF-α, IL-6, and IL-17) in ovx mice, which were significantly reduced by both V1 and V2 treatments **(Figure 2G).** In contrast, anti-osteoclastogenic cytokines (IFN-γ, TGF-β, and IL-10) were suppressed in ovx mice but restored upon V1 and V2 supplementation **(Figure 2F).** Collectively, these results demonstrate that vitamin D supplementation mitigates bone loss and improves bone quality in ovx mice, with both doses exerting pronounced protective effects on bone microarchitecture, mechanical strength, and osteoimmune regulation.

### 3.2 Vitamin D promotes osteoclastogenesis in an ILC3-mediated manner

We next investigated the mechanisms underlying vitamin D-mediated inhibition of bone resorption. To determine whether vitamin D directly affects osteoclast differentiation, BMCs were cultured in osteoclastogenic medium containing M-CSF (30 ng/mL) and RANKL (60 ng/mL), with or without vitamin D (50 nM). Interestingly, vitamin D markedly enhanced osteoclastogenesis, as evidenced by an increased number of TRAP-positive cells and cells with more than 3 nuclei **(Figures 3A-C).** Moreover, the osteoclast area is significantly increased in the vitamin D-treated group compared to the control group **(Figure 3D).** These findings suggest that vitamin D directly promotes osteoclast formation, implying that its protective effects on bone health are likely mediated through indirect mechanisms. Consistent with this notion, *in vivo* serum cytokine analysis revealed that vitamin D downregulated osteoclastogenic cytokines while upregulating anti-osteoclastogenic cytokines **(Figure 2G),** suggesting an immunomodulatory mechanism underlying its bone-protective effects. Since vitamin D exerts profound immunomodulatory effects on both innate and adaptive immune cells, we first investigated its role in maintaining bone health via modulating innate immune cells. Given previous evidence that ILCs (particularly ILC3) are regulated by vitamin D, we first examined whether vitamin D influences ILC populations in the BM to maintain bone homeostasis. We observed that the frequency of the ILC1 and ILC2 cells remains unaffected in all the groups **(Figures 3E-H).** However, the frequency of ILC3s was significantly reduced in ovx mice, whereas administration of V1 and V2 did not alter the overall ILC3 frequency **(Figures 3I-J).** Since ILC3s comprise IL-17□ and IL-22□ subsets, we further investigated the effects of vitamin D on these populations. The IL-17□ ILC3 population was markedly elevated in ovx mice but remained unchanged following vitamin D administration at either concentration **(Figure 3K).** In contrast, while the frequency of IL-22□ ILC3s was comparable between sham and ovx groups, V2 administration significantly enhanced IL-22 secretion **(Figure 3L).** We next evaluated the effect of vitamin D on ILC3s under *in vitro* conditions. Immune cells isolated from the spleen were subjected to magnetic separation to obtain lineage-negative cells using a biotin-labeled antibody cocktail (anti-CD3, CD19, F4/80, NK1.1, FcεRI, Gr-1, CD5, CD11c, TER119). The negatively selected cells were then cultured under ILC3-polarizing conditions (IL-2, IL-7, IL-23, and IL-1β) in the presence or absence of vitamin D (50 nM) for 24, 48, and 72 hours. Time-kinetic analysis revealed that vitamin D did not affect the overall frequency of ILC3s at any time point **(Figures 4A-B)**. However, at 24 hours, vitamin D significantly modulated cytokine production, decreasing IL-17 while increasing IL-22 levels **(Figures 4C-F).** Based on these findings, 24 hours was identified as the optimal treatment duration for subsequent functional assays. To assess the impact of ILC3s on osteoclast differentiation, BMCs were co-cultured with ILC3s. Interestingly, it was observed that unprimed ILC3s suppressed osteoclast formation in a cell ratio-dependent manner **(Figures 4G-J).** However, in contrast, vitamin D-primed ILC3s promoted osteoclastogenesis, increasing both the number and area of TRAP-positive multinucleated cells **(Figures 4G-J).** Collectively, these results indicate that vitamin D promotes osteoclastogenesis through both direct and indirect mechanisms, the latter mediated by vitamin D-induced ILC3s. Since vitamin D administration enhances IL-22 secretion from ILC3s, we hypothesized that vitamin D-primed ILC3s might influence bone remodeling through IL-22. To test this, we examined the direct effect of IL-22 on osteoclastogenesis of the BMCs *in vitro* using varying concentrations of recombinant IL-22. Notably, treatment with IL-22 at a concentration of 10 ng/ml significantly increased the osteoclast area **(Supplementary Figure 6A-C)**, suggesting that elevated IL-22 secretion from vitamin D-primed ILC3s might contribute to enhanced osteoclastogenesis. To further investigate the mechanistic role of IL-22 in ILC3-mediated osteoclastogenesis, we performed *in vitro* co-culture experiments using BMCs and vitamin D-primed ILC3s in the presence of a neutralizing α-IL-22 antibody. Notably, blockade of IL-22 significantly reduced osteoclast differentiation compared to control conditions, as indicated by a reduced number of trap-positive cells with more than 3 nuclei and area of osteoclasts in the α-IL-22-treated group **(Figure 4J-L)**. These findings demonstrate that IL-22 is a critical mediator of the pro-osteoclastogenic effect of vitamin D-primed ILC3s and establish a functional link between ILC3-derived IL-22 and enhanced osteoclastogenesis.

**Fig. 3:**
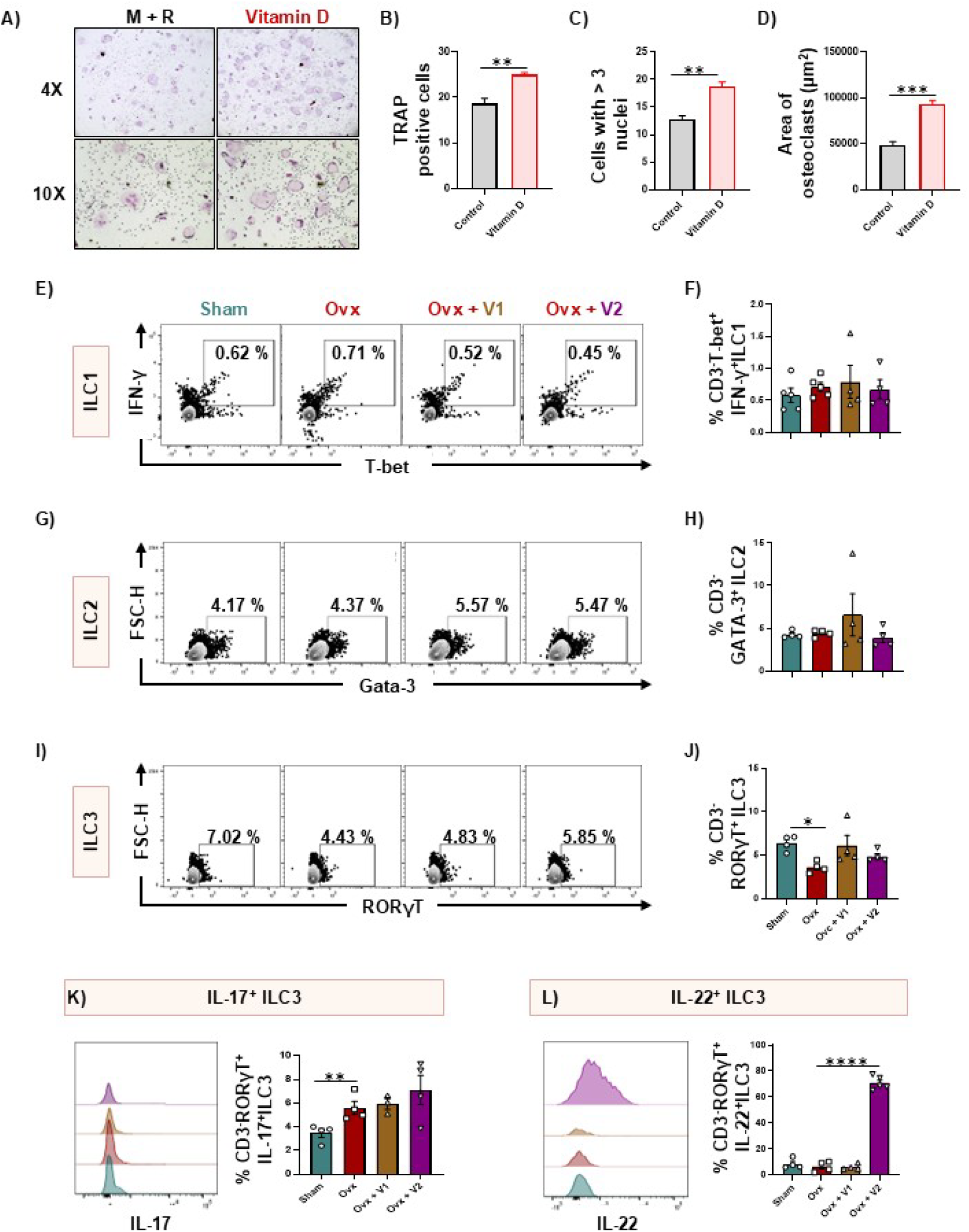
Vitamin D promote osteoclastogenesis of bone marrow cells (BMCs). BMCs were harvested from mice and were cultured in the presence or absence of vitamin D (50 nM) along with MCSF (30 ng/ml) and RANKL (60 ng/ml). **(A)** Pictograph images (4X and 10X) representing osteoclastogenesis in the presence and absence of vitamin D. **(B)** Bar graph representing the number of TRAP-positive cells. **(C)** Bar graph representing the number of multinucleated cells with more than three nuclei. **(D)** Bar graph representing the area of a multinucleated osteoclast. Cells from the bone marrow (BM) of all groups were harvested and analysed by flow cytometry to determine the percentage of ILCs. **(E & F)** Contour plot and bar graph depicting percentages of ILC1 (CD3^-^T-bet^+^IFN-γ^+^) in BM of sham, ovx, ovx + V1 and ovx + V2 groups. **(G & H)** Contour plot and bar graph depicting percentages of ILC2 (CD3^-^GATA-3^+^) in BM of sham, ovx, ovx +V1 and ovx + V2 groups. **(I & J)** Contour plot and bar graph depicting percentages of ILC3 (CD3^-^RORγT^+^) in BM of sham, ovx, ovx + V1 and ovx + V2 groups. **(K)** Overlay plot and bar graph depicting percentages of IL-17^+^ ILC3 (CD3^-^RORγT^+^IL-17^+^) in BM of sham, ovx, ovx + V1 and ovx + V2 groups. **(L)** Overlay plot and bar graph depicting percentages of IL-22^+^ ILC3 (CD3^-^RORγT^+^IL-22^+^) in BM of sham, ovx, ovx +V1 and ovx + V2 groups. Data are represented as Mean ± SEM. The results were evaluated by one-way ANOVA followed by Dunnett’s. Statistical significance was defined as *p < 0.05, **p < 0.01, ***p < 0.001 and ****p < 0.0001, concerning the indicated mouse group.

**Fig. 4:**
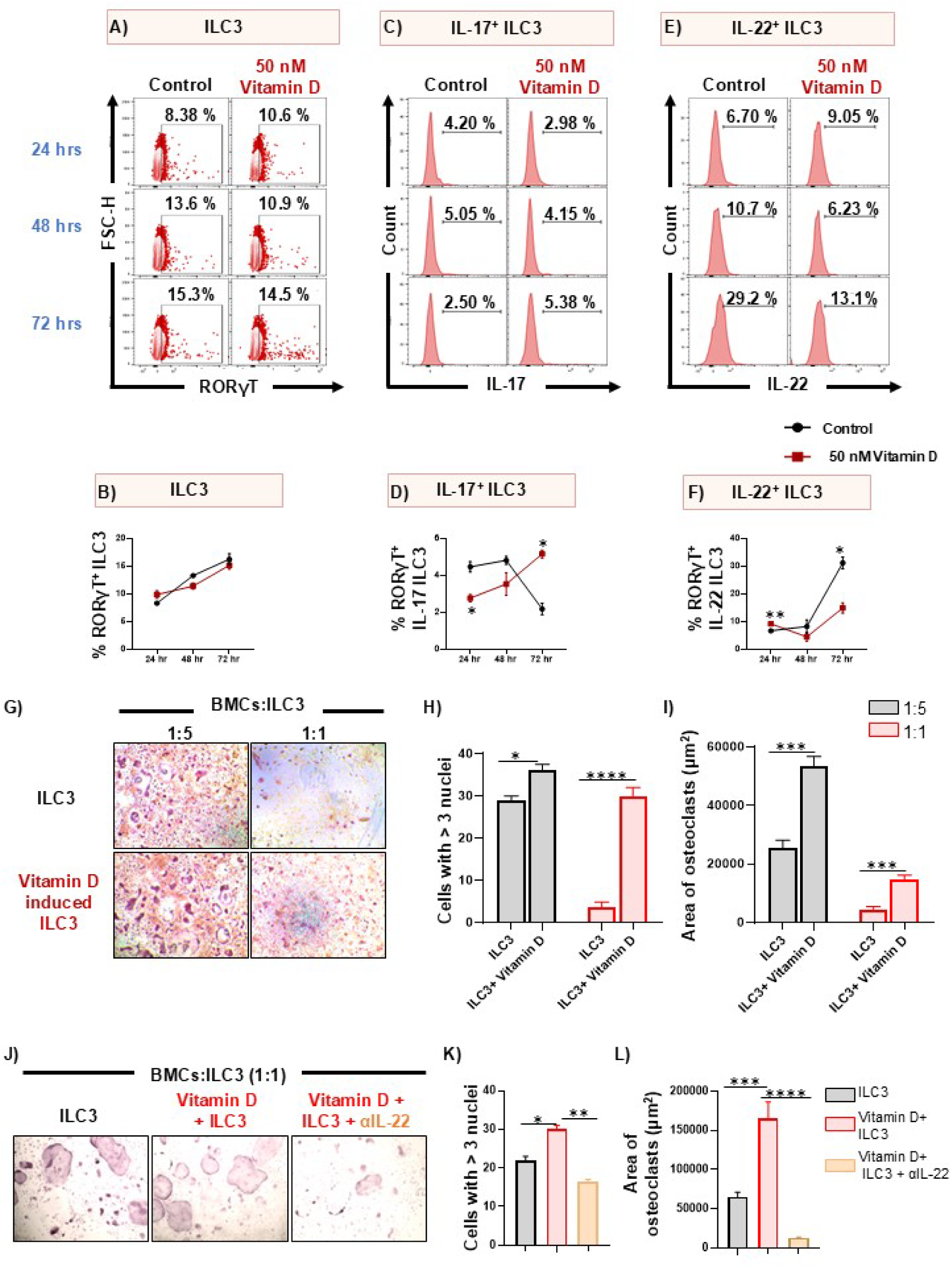
Vitamin D-primed ILC3 promote osteoclastogenesis of bone marrow cells (BMCs). The spleen was harvested and processed for the negative selection of ILCs, followed by stimulation with IL-2 (10 ng/ml), IL-7 (10 ng/ml), IL-1β (50 ng/ml), and IL-23 (50 ng/ml) for 24, 48 and 72 hours and flow cytometry was performed to analyze the population of **(A & B)** ILC3s, **(C & D)** IL-17^+^ ILC3s, and **(E & F)** IL-22^+^ ILC3s. BMCs were harvested from mice and were co-cultured with ILC3s stimulated in the presence and absence of vitamin D for 24 hours, along with MCSF (30 ng/ml) and RANKL (60 ng/ml). **(G)** Pictograph images (10X) representing osteoclastogenesis in the presence of vitamin D, unprimed and primed ILC3s. **(H)** Bar graph representing the number of multinucleated cells with more than three nuclei. **(I)** Bar graph representing the area of a multinucleated osteoclast. BMCs and vitamin D-induced ILC3s were co-cultured (1:1) in media supplemented with M-CSF (30 ng/ml) and RANKL (60 ng/ml), along with the presence or absence of α-IL-22 (10 μg/ml) for 4 days. **(J)** Pictographic images (10X) depicting osteoclastogenesis under different conditions: unprimed ILC3s, vitamin D-primed ILC3s, and vitamin D-primed ILC3s in the presence of α-IL-22. **(K)** Bar graph representing the number of multinucleated cells with more than three nuclei. **(L)** Bar graph representing the area of a multinucleated osteoclast. Data are represented as Mean ± SEM. The results were evaluated by one-way ANOVA followed by Dunnett’s test. Statistical significance was defined as *p < 0.05, **p < 0.01, ***p < 0.001 and ****p < 0.0001, concerning the indicated mouse group.

### 3.3 Vitamin D inhibit osteoclastogenesis in a Treg-mediated manner

Since vitamin D appeared to protect bone health not in an ILC3-mediated manner, we next investigated whether its effects were mediated through T cells. To evaluate this, naïve T cells were magnetically selected and cultured in the presence or absence of vitamin D (50 nM) for five days. After induction, these vitamin D-treated T cells were co-cultured with BMCs in osteoclastogenic medium containing M-CSF and RANKL. Interestingly, vitamin D-induced T cells significantly inhibited osteoclast differentiation, as indicated by a marked reduction in the number of TRAP-positive multinucleated cells (with ≥3 nuclei) and a decrease in osteoclast area at a 1:1 ratio **(Figures 5A-C).** These findings suggest that vitamin D may exert anti-osteoclastogenic effects indirectly by enhancing the regulatory potential of T cells. Given that T cells consist of distinct functional subsets, we next examined which subsets are modulated by vitamin D *in vivo*. We analysed the frequencies of Th1, Th2, and Th17 cells in sham and ovx mice treated with vitamin D (V1 and V2). The frequencies of IFNγ□ Th1 cells were significantly elevated, whereas Th2 cell frequencies were markedly reduced in ovx mice compared with sham controls; however, vitamin D treatment at both concentrations did not alter either population, indicating that its effect on bone health is not Th1– or Th2-dependent **(Figures 5D-I).** We then focused on Th17 cells, which are known to play a pivotal role in bone remodeling. As reported previously, ovx mice displayed elevated Th17 cell frequency and increased IL-17 secretion compared with sham controls **(Figures 6A-C).** V2 administration significantly reduced Th17 cell frequency in ovx mice, and concurrently reduced IL-17 secretion by these cells **(Figures 6A-C).** Since IL-22 can also be produced by Th17 cells, we next analyzed IL-22□ Th17 cells across groups. Interestingly, while no significant difference was observed in the frequency of IL-22□ Th17 cells between sham and ovx mice, V2 administration markedly increased IL-22□ Th17 cells, suggesting that higher concentrations of vitamin D preferentially enhance IL-22 production from both ILC3s and Th17 cells **(Figure 6D).** To further explore the impact of vitamin D on Th17-mediated osteoclastogenesis, naïve T cells were cultured under Th17-polarising conditions in the presence or absence of vitamin D. Vitamin D treatment significantly reduced Th17 cell differentiation while simultaneously augmenting IL-22 secretion from these cells, thereby corroborating the *in vivo* findings **(Supplementary Figure 7A-D)**. Vitamin D-primed Th17 cells were then further co-cultured with BMCs in the presence of MCSF and RANKL. Notably, we observed that compared with unprimed Th17 cells, vitamin D-primed Th17 cells further enhanced osteoclast formation, as indicated by an increased number of TRAP-positive multinucleated cells, without significant changes in osteoclast area **(Figures 6E-G).** These results suggest that vitamin D enhances the osteoclastogenic potential of Th17 cells, possibly through upregulation of IL-22 production. Next, we examined whether there was any correlation between Th17 cell frequency and serum vitamin D levels in human PMO subjects. For this, CD4□IL-17□ Th17 cells were analysed in PBMCs by flow cytometry. Our results demonstrated a negative correlation between Th17 cell frequency and serum vitamin D levels in PMO subjects **(Figure 6H).** In addition, serum IL-17 levels were also inversely correlated with vitamin D concentration **(Figure 6I),** suggesting that vitamin D may exert suppressive effects on Th17 cells.

**Fig. 5:**
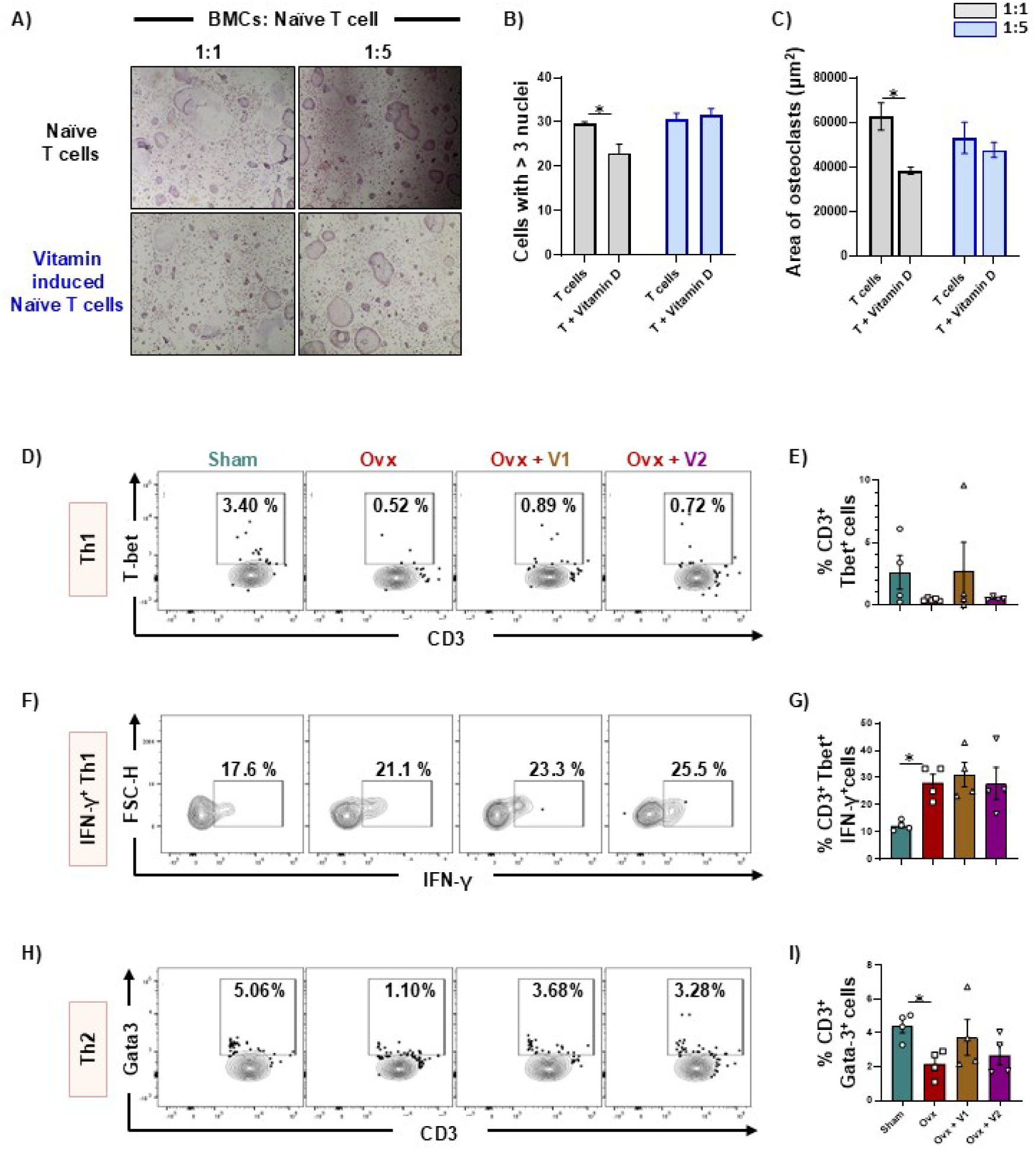
Vitamin D inhibits osteoclastogenesis in a T cell-dependent manner. The spleen was harvested and processed for the magnetic selection of naïve T cells, which were then stimulated with vitamin D (50 nM) for 5 days. Bone marrow cells (BMCs) were harvested from mice and were co-cultured with naïve T cells stimulated in the presence and absence of vitamin D, along with MCSF (30 ng/ml) and RANKL (60 ng/ml) for 4 days. **(A)** Pictograph images (10X) representing osteoclastogenesis in the presence of vitamin D-unprimed and vitamin D-primed naïve T cells. **(B)** Bar graph representing the number of multinucleated cells with more than three nuclei. **(C)** Bar graph representing the area of a multinucleated osteoclast. **(D & E)** Contour plot and bar graph depicting percentages of Th1 (CD3^+^Tbet^+^) cells in the bone marrow (BM) of sham, ovx, ovx + V1 and ovx + V2 groups. **(F & G)** Contour plot and bar graph depicting percentages of IFN-γ^+^ Th1 cells (CD3^+^Tbet^+^IFN-γ^+^) in BM of sham, ovx, ovx +V1 and ovx + V2 groups. **(H & I)** Contour plot and bar graph depicting percentages of Th2 cells (CD3^+^GATA-3^+^) in the BM of sham, ovx, ovx + V1 and ovx + V2 groups. Data are represented as Mean ± SEM. The results were evaluated by one-way ANOVA followed by Dunnett’s test. Statistical significance was defined as *p < 0.05, **p < 0.01, ***p < 0.001 and ****p < 0.0001, concerning the indicated mouse group.

**Fig. 6:**
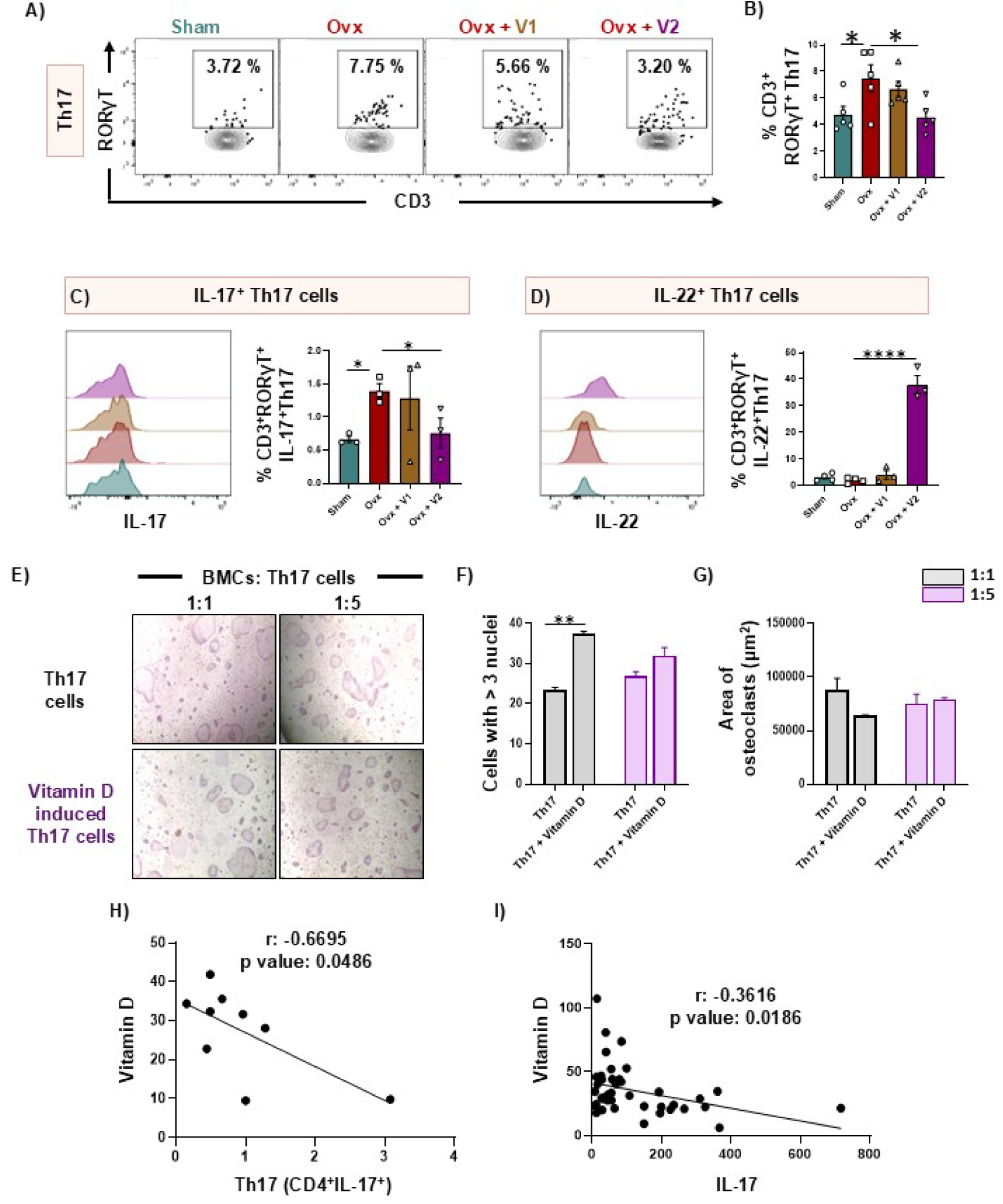
Vitamin D-primed Th17 cells have more potent osteoclastogenic potential. Cells from the bone marrow (BM) of all the groups were harvested and analyzed by flow cytometry for the percentage of Th17 cells (CD3^+^ROR_γ_T^+^). **(A & B)** Contour plot and bar graph depicting percentages of Th17 in BM of sham, ovx, ovx + V1 and ovx + V2 groups. **(C)** Overlay plot and bar graph depicting percentages of IL-17^+^ Th17 (CD3^+^ROR_γ_T^+^IL-17^+^) in BM of sham, ovx, ovx + V1 and ovx + V2 groups. **(D)** Overlay plot and bar graph depicting percentages of IL-22^+^ Th17 (CD3^+^ROR_γ_T^+^IL-22^+^) in BM of sham, ovx, ovx +V1 and ovx + V2 groups. Bone marrow cells (BMCs) were harvested from mice and were co-cultured with Th17 stimulated in the presence and absence of vitamin D for 5 days, along with MCSF (30 ng/ml) and RANKL (60 ng/ml). **(E)** Pictograph images (10X) representing osteoclastogenesis in the presence of vitamin D-unprimed and vitamin D-primed Th17 cells. **(F)** Bar graph representing the number of multinucleated cells with more than three nuclei. **(G)** Bar graph representing the area of a multinucleated osteoclast. **(H)** Correlation between Th17 cells (PBMCs) and vitamin D levels in the human subjects. **(I)** Correlation between IL-17 cytokine (serum) and vitamin D levels in the human subjects. Data are represented as Mean ± SEM. The results were evaluated by one-way ANOVA followed by Dunnett’s test. Statistical significance was defined as *p < 0.05, **p < 0.01, ***p < 0.001 and ****p < 0.0001, concerning the indicated mouse group.

Another key T cell subset with a critical role in maintaining bone homeostasis is the Treg population. Therefore, we next analysed the status of Tregs following vitamin D administration in ovx mice. As reported previously, both total Tregs and IL-10□ Tregs were significantly reduced in ovx mice compared with sham controls **(Figures 7A-D).** Interestingly, V1 supplementation markedly increased the overall Treg population, while V2 supplementation significantly enhanced the frequency of IL-10□ Tregs **(Figures 7A-D).** Serum cytokine analysis further revealed elevated levels of TGF-β and IL-10, key cytokines associated with Tregs in both the V1 and V2 treatment groups **(Figure 2G).** These findings suggest that vitamin D may exert its bone-protective effects, at least in part, through the modulation and activation of Tregs. To further assess the functional impact of vitamin D-induced Tregs on osteoclastogenesis, naïve T cells were magnetically isolated and cultured under Treg-polarizing conditions for five days in the presence or absence of vitamin D. Vitamin D was observed to significantly promote Treg differentiation, accompanied by a marked increase in IL-10 secretion **(Supplementary Figure 8A-D).** Vitamin D-induced Tregs were then further co-cultured with BMCs under osteoclastogenic conditions, as described earlier. Remarkably, vitamin D-induced Tregs exhibited a stronger inhibitory effect on osteoclast differentiation, as evidenced by a significant reduction in both the number of TRAP-positive multinucleated cells (≥3 nuclei) and the osteoclast area **(Figures 7E-G).** Furthermore, we analyzed the CD4□IL-10□ T cell population within PBMCs of human subjects and observed a positive correlation between these cells and serum vitamin D levels **(Figure 7H).** In contrast, total serum IL-10 concentrations showed no significant correlation with vitamin D levels, suggesting that vitamin D specifically enhances IL-10 production by CD4□ T cells rather than broadly increasing systemic IL-10 levels (**Figure 7I)**.

**Fig. 7:**
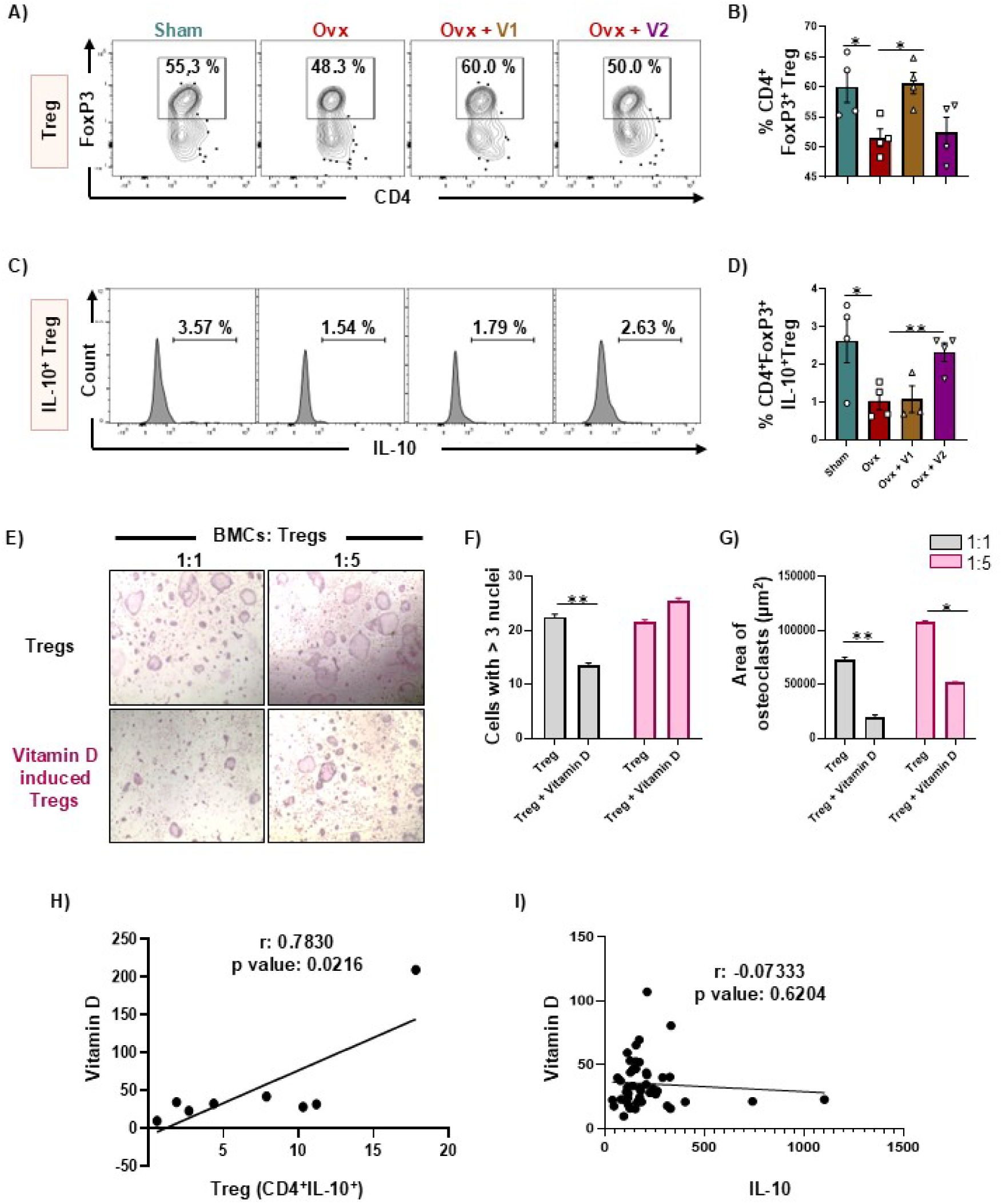
Vitamin D enhances the anti-osteoclastogenic potential of Tregs. Cells from the bone marrow (BM) of all the groups were harvested and analyzed by flow cytometry for the percentage of Tregs (CD4^+^Foxp3^+^). **(A & B)** Contour plot and bar graph depicting percentages of Tregs in the BM of sham, ovx, ovx + V1 and ovx + V2 groups. **(C & D)** Contour plot and bar graph depicting percentages of IL-10^+^ Tregs (CD3^+^ROR_γ_T^+^IL-10^+^) in BM of sham, ovx, ovx + V1 and ovx + V2 groups. Bone marrow cells (BMCs) were harvested from mice and co-cultured with Tregs stimulated in the presence and absence of vitamin D for 5 days, along with MCSF (30 ng/ml) and RANKL (60 ng/ml). **(E)** Pictograph images (10X) representing osteoclastogenesis in the presence of vitamin D, unprimed and primed Tregs. **(F)** Bar graph representing the number of multinucleated cells with more than three nuclei. **(G)** Bar graph representing the area of a multinucleated osteoclast. **(H)** Correlation between Treg cells (PBMCs) and vitamin D levels in the human subjects. **(I)** Correlation between IL-10 cytokine (serum) and vitamin D levels in the human subjects. Data are represented as Mean ± SEM. The results were evaluated by one-way ANOVA followed by Dunnett’s test. Statistical significance was defined as *p < 0.05, **p < 0.01, ***p < 0.001 and ****p < 0.0001, concerning the indicated mouse group.

Together, these results indicate that vitamin D induce bone loss in an ILC3-Th17-mediated manner but protects bone health by enhancing the anti-osteoclastogenic potential of Tregs, thereby contributing to the suppression of bone resorption and maintenance of bone homeostasis.

### 3.4 Vitamin D suppress osteoclast differentiation in a Breg-dependent manner

Treg and Th17 cell balance is further regulated by another very crucial adaptive regulatory immune cell, i.e. Breg. Recent studies from our group have demonstrated that Bregs mitigate bone loss in PMO by inhibiting osteoclastogenesis and modulating the Treg-Th17 cell balance. Building on this, we next examined whether Bregs contribute to the bone-protective effects of vitamin D. Consistent with our previous findings, both the frequency of total Bregs and IL-10□ Bregs were markedly reduced in ovx mice compared to sham controls **(Figures 8A-D).** Interestingly, treatment with both V1 and V2 significantly increased the proportion of Bregs and IL-10□ Bregs, suggesting that vitamin D preferentially influences B cell populations over other immune cells, and that its bone-protective effects may be mediated, at least in part, through Bregs **(Figures 8A-D).** Serum cytokine analysis further revealed elevated levels of TGF-β and IL-10– key cytokines associated with Tregs in both the V1 and V2 treatment groups **(Figure 2G).**

**Fig. 8:**
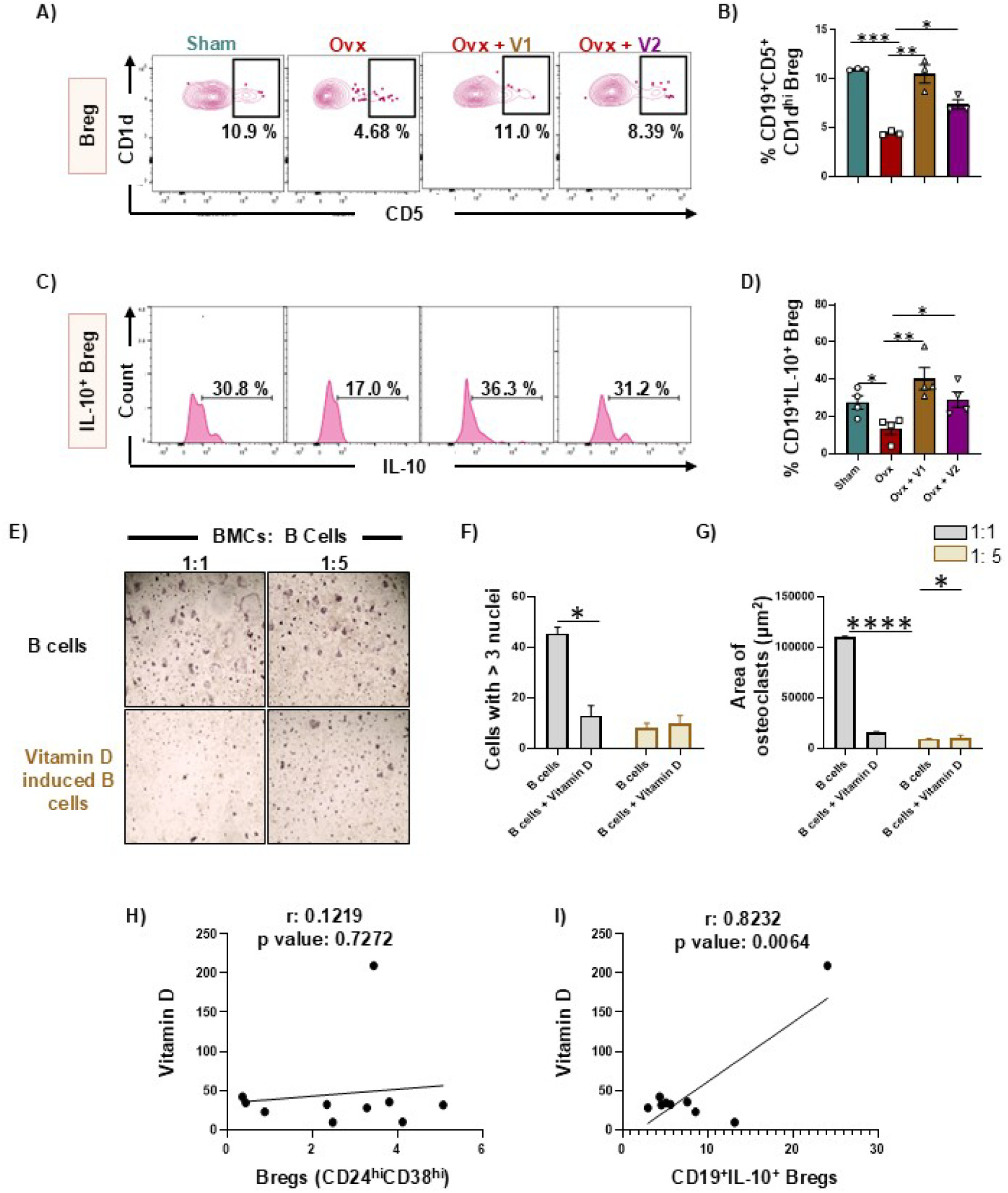
Vitamin D enhances the anti-osteoclastogenic potential of Bregs. Cells from the bone marrow (BM) of all the groups were harvested and analyzed by flow cytometry for the percentage of Bregs (CD19^+^CD5^+^CD1d^hi^). **(A & B)** Contour plot and bar graph depicting percentages of Bregs in BM of sham, ovx, ovx + V1 and ovx + V2 groups. **(C & D)** Contour plot and bar graph depicting percentages of IL-10^+^ Bregs (CD19^+^IL-10^+^) in BM of sham, ovx, ovx + V1 and ovx + V2 groups. Bone marrow cells (BMCs) were harvested from mice and co-cultured with B cells stimulated with vitamin D for 24 hours, along with MCSF (30 ng/ml) and RANKL (60 ng/ml). **(E)** Pictograph images (10X) representing osteoclastogenesis in the presence of vitamin D, unprimed and primed Bregs. **(F)** Bar graph representing the number of multinucleated cells with more than three nuclei. **(G)** Bar graph representing the area of a multinucleated osteoclast. The results were evaluated using the unpaired Student t-test. **(H)** Correlation between Breg cells (CD38^hi^CD24^hi^) and vitamin D levels in the human subjects. **(I)** Correlation between Breg cells (CD19^+^IL-10^+^) and vitamin D levels in the human subjects. Data are represented as Mean ± SEM. The results were evaluated by one-way ANOVA followed by Dunnett’s test. Statistical significance was defined as *p < 0.05, **p < 0.01, ***p < 0.001 and ****p < 0.0001, concerning the indicated mouse group.

To further investigate this, we magnetically selected B cells and stimulated them with vitamin D for 24 hours to induce Breg differentiation. Vitamin D treatment was observed to significantly enhance the differentiation of Bregs (B220□ IL-10□) **(Supplementary Figure 9A-B).** Vitamin D-induced Bregs were then co-cultured with BMCs as described above. We observed that Bregs markedly inhibited osteoclastogenesis in BMCs **(Figures 8E-G)**, highlighting a key role for vitamin D in promoting bone health by enhancing the anti-osteoclastogenic potential of Bregs. Further, we analyzed the Breg cell population in PBMCs of human PMO subjects and observed that CD19□IL-10□ Bregs **(Figures 8H-I)** exhibited a positive correlation with serum vitamin D levels, underscoring the pivotal role of vitamin D in inducing Breg differentiation. Moreover, since Bregs act upstream of Tregs and Th17 cells and play a regulatory role in their differentiation, our findings suggest that vitamin D may mitigate bone loss primarily through a Breg-mediated immunomodulatory mechanism.

### 3.5 Vitamin D prevents bone loss by maintaining gut permeability

We observed that the expression of enzymes involved in the synthesis and regulation of active vitamin D, namely 1α-hydroxylase and 24-hydroxylase, was markedly reduced in the colon of ovx mice **(Figure 9A-B).** Notably, vitamin D supplementation (V2) restored the expression of both enzymes in colonic tissue, indicating impaired local synthesis of the active form of vitamin D in ovx mice **(Figure 9A-B).** Additionally, vitamin D receptor (VDR) expression in the colon was significantly diminished in ovx mice but was enhanced following vitamin D (V2) administration **(Figure 9C).** Collectively, these findings suggest a critical role for vitamin D signaling in the colon, prompting us to further investigate the functional impact of vitamin D in colonic tissue. Our previous data indicated that vitamin D markedly enhances IL-22 production from both ILC3s and Th17 cells. Given that IL-22 plays a critical role in maintaining gut barrier integrity-and that increased gut permeability has been closely linked to bone loss-we next investigated whether vitamin D protects bone health by modulating IL-22 production from ILC3s and Th17 cells in the MLNs and colon. We observed a significant reduction in the frequency of ILC3s in both the MLNs and colon of ovx mice compared to sham controls **(Figures 9D-E).** Notably, vitamin D treatment did not affect the total ILC3 population or IL-17 secretion from these cells **(Figures 9F-G).** However, the proportion of IL-22□ ILC3s was markedly decreased in ovx mice, and V1 and V2 administration significantly restored IL-22□ ILC3 levels in both the MLNs and colon **(Figures 9H-I).** Next, we examined the effect of vitamin D on gut Th17 cells. Vitamin D administration significantly reduced the frequency of Th17 cells in both the MLNs and colon, although it did not significantly alter IL-17 or IL-22 secretion from this subset **(Figures 10A-F).** Furthermore, assessment of gut permeability revealed a pronounced increase in intestinal leakage in ovx mice, which was significantly reversed following vitamin D treatment at both concentrations **(Figure 11A).** The vitamin D-dependent prevention of increased gut permeability can be explained by its ability to enhance the expression of claudin and IL-22 genes while concomitantly reducing IL-17 levels in colonic tissue **(Figure 11B-E).**

**Fig. 9:**
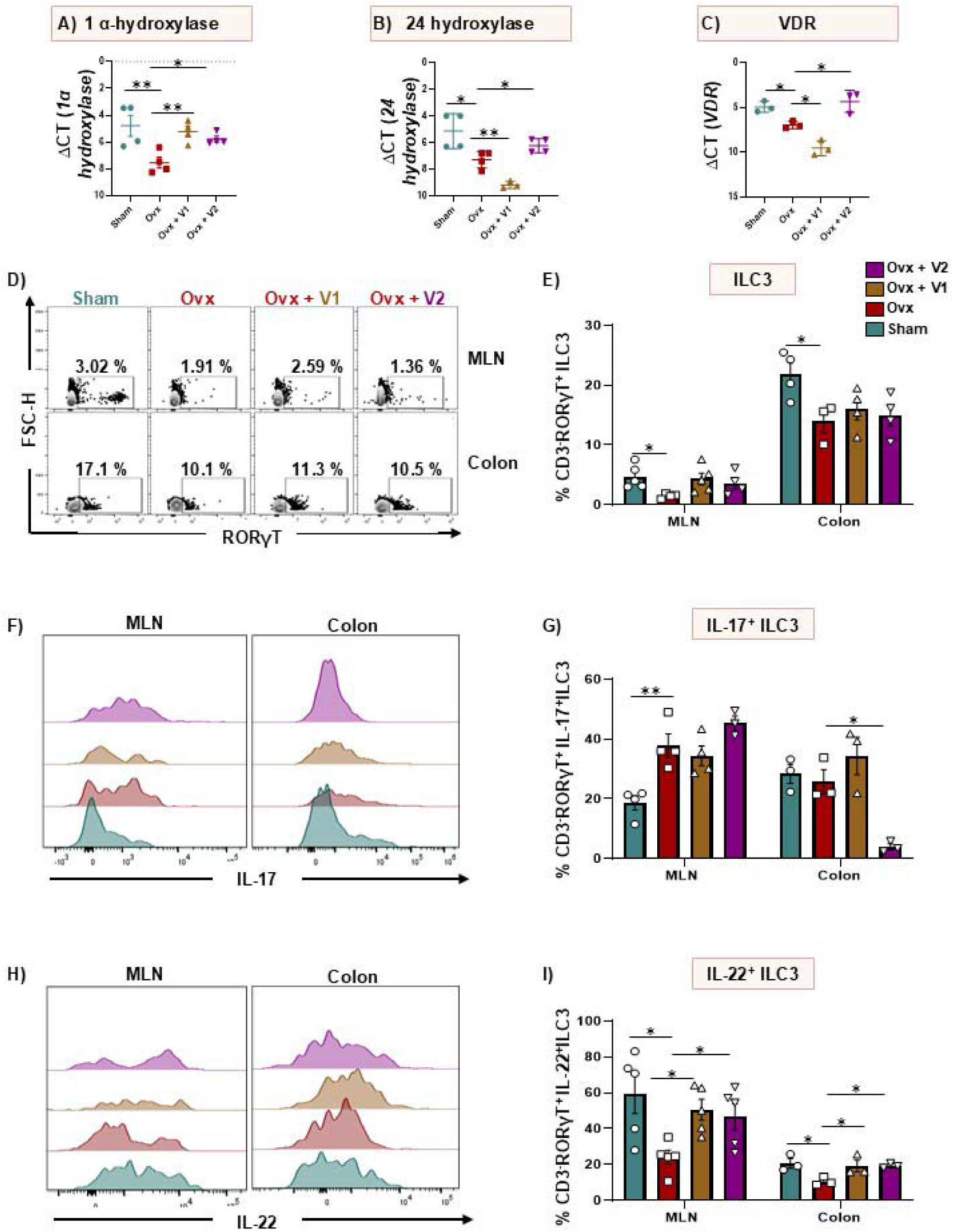
Vitamin D administration enhances IL-22 production from ILC3s. Gene expression analysis of **(A)** 1 _α_-hydroxylase, **(B)** 24-hydroxylase, and **(C)** vitamin D receptor (VDR) in the colon of all groups of mice. Cells from the mesenteric lymph node (MLN) and colon of all the groups were harvested and analyzed by flow cytometry for the percentage of ILC3 (CD3^-^ROR_γ_T^+^). **(D & E)** Contour plot and bar graph depicting percentages of ILC3 in MLN and colon of sham, ovx, ovx + V1 and ovx + V2 groups. **(F & G)** Overlay plot and bar graph depicting percentages of IL-17^+^ ILC3 (CD3^-^ROR_γ_T^+^IL-17^+^) in MLN and colon of sham, ovx, ovx + V1 and ovx + V2 groups. **(H & I)** Overlay plot and bar graph depicting percentages of IL-22^+^ ILC3 (CD3^-^ROR_γ_T^+^IL-22^+^) in MLN and colon of sham, ovx, ovx + V1 and ovx + V2 groups. Data are represented as Mean ± SEM. The results were evaluated by one-way ANOVA followed by Dunnett’s test. Statistical significance was defined as *p < 0.05, **p < 0.01, ***p < 0.001 and ****p < 0.0001, concerning the indicated mouse group.

**Fig. 10:**
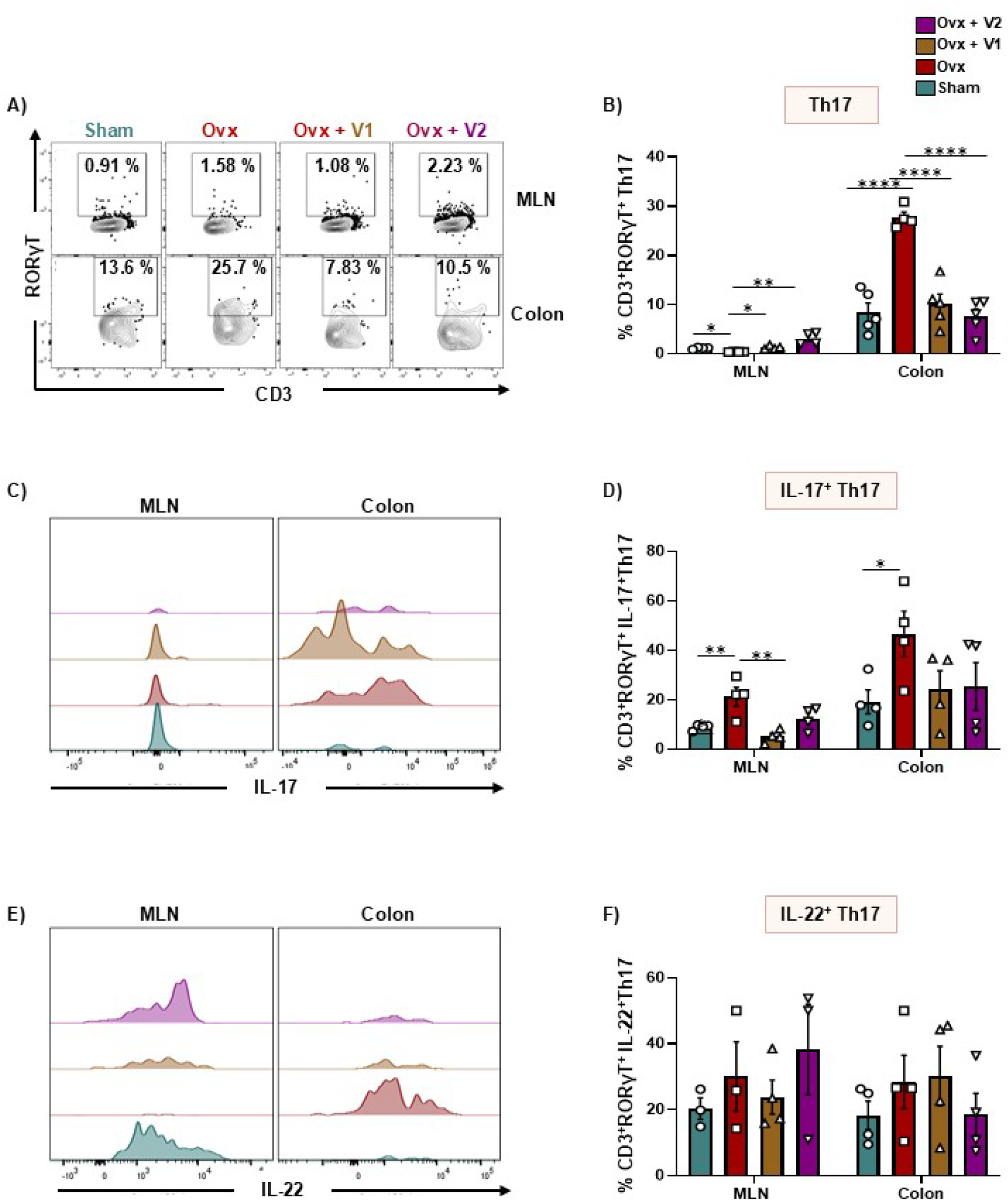
Vitamin D administration does not influence IL-22 production from Th17 cells. Cells from the mesenteric lymph nodes (MLN) and colon of all the groups were harvested and analyzed by flow cytometry for the percentage of Th17 (CD3^+^ROR_γ_T^+^) cells. **(A & B)** Contour plot and bar graph depicting percentages of Th17 in MLN and colon of sham, ovx, ovx + V1 and ovx + V2 groups. **(C & D)** Overlay plot and bar graph depicting percentages of IL-17^+^ Th17 (CD3^+^ROR_γ_T^+^IL-17^+^) in MLN and colon of sham, ovx, ovx + V1 and ovx + V2 groups. **(E & F)** Overlay plot and bar graph depicting percentages of IL-22^+^ Th17 (CD3^+^ROR_γ_T^+^IL-22^+^) in MLN and colon of sham, ovx, ovx + V1 and ovx + V2 groups. Data are represented as Mean ± SEM. The results were evaluated by one-way ANOVA followed by Dunnett’s test. Statistical significance was defined as *p < 0.05, **p < 0.01, ***p < 0.001 and ****p < 0.0001, concerning the indicated mouse group.

**Fig. 11:**
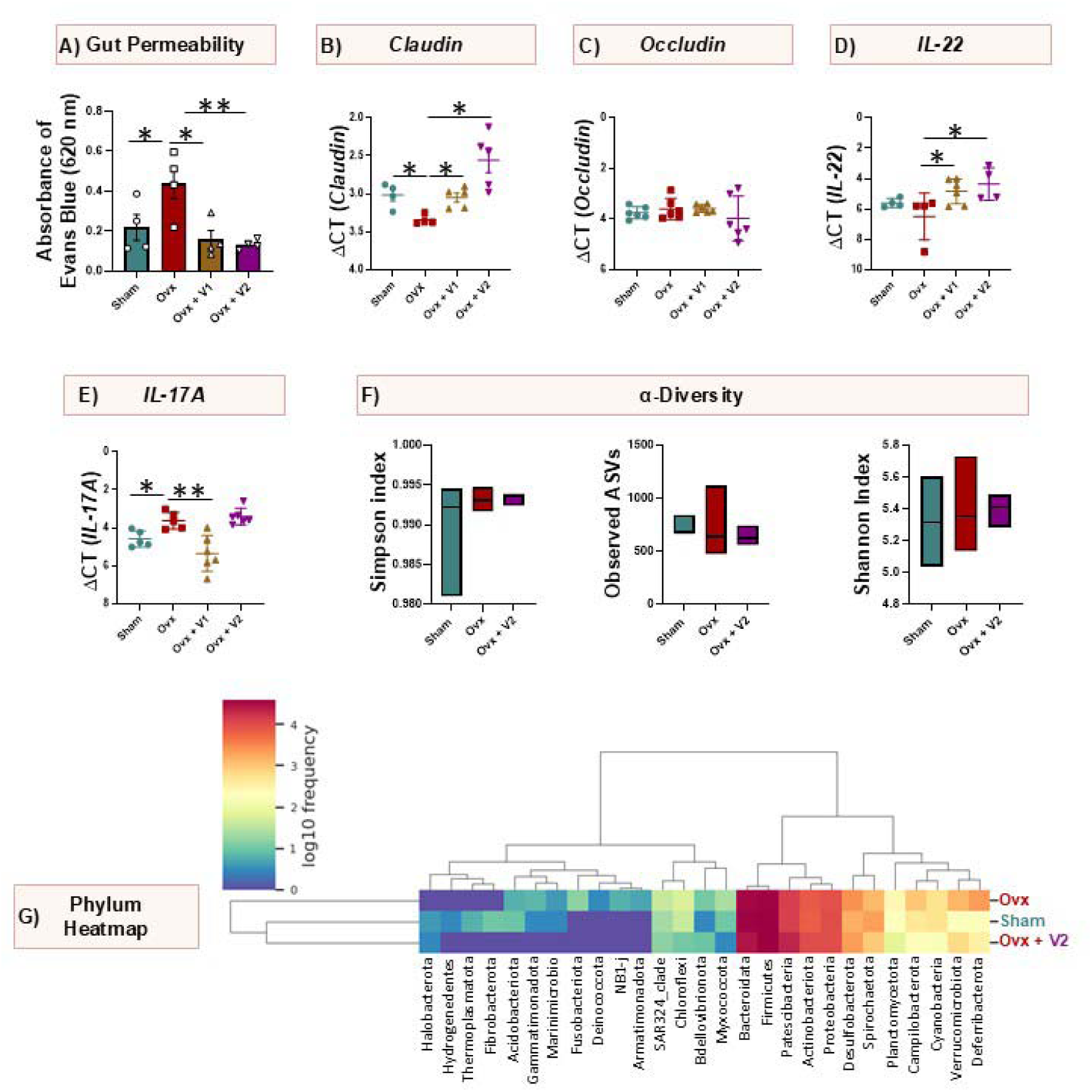
Vitamin D administration prevents dysbiosis in Ovx mice: **(A)** Measurement of gut integrity. Gene expression analysis of **(B)** claudin, **(C)** occludin, **(D)** IL-22, and **(E)** IL-17. **(F)** Bar graphs representing alpha diversity (Simpson index, Observed ASVs and Shannon index) in each group. **(G)** Heatmap denoting the dominant phylum in each group. Data are represented as Mean ± SEM. The results were evaluated by one-way ANOVA followed by Dunnett’s test. Statistical significance was defined as *p < 0.05, **p < 0.01, ***p < 0.001 and ****p < 0.0001, concerning the indicated mouse group.

Collectively, these findings suggest that vitamin D preserves gut barrier integrity by enhancing IL-22 production from ILC3s. This mechanism likely contributes to the bone-protective effects of vitamin D, linking gut immune regulation to skeletal homeostasis.

### 3.6 Vitamin D restores bone health by restoring gut microbial composition

Increased intestinal permeability is closely associated with bone loss, and previous studies, including ours, have shown that gut dysbiosis exacerbates bone deterioration. To explore whether vitamin D (V2) supplementation could modulate gut dysbiosis, we conducted a comprehensive gut microbiome analysis across all experimental groups. Our results revealed a reduction in Simpson index, Observed ASVs, and Shannon index in the ovx + V2 group, suggesting decreased microbial diversity following vitamin D treatment **(Figure 11F).** At the phylum level, Firmicutes and Bacteroidetes remained dominant across all groups. Notably, the abundance of Desulfobacterota, a phylum linked to increased gut permeability, was elevated in ovx mice but reduced following V2 administration **(Figures 11G)**. In contrast, vitamin D treatment enhanced the relative abundance of Actinobacteriota and Patescibacteria, phyla often associated with gut health and metabolic balance **(Figures 11G)**. At the genus level, the beneficial *Lactobacillus* population, markedly diminished in ovx mice, was effectively restored by V2 supplementation **(Figures 12A-B)**. Furthermore, vitamin D increased the abundance of *Clostridium UCG-014*, known producers of SCFAs that play a crucial role in maintaining intestinal integrity and supporting bone metabolism **(Figures 12A-B)**. As SCFA-producing bacteria were reduced in ovx mice but restored upon V2 administration, we next examined whether SCFAs influence vitamin D-mediated osteoclastogenesis. For this, BMCs were cultured with vitamin D in the presence or absence of an SCFA mixture (0.5 mM each of acetate, propionate, butyrate, and valerate, as determined from our previous findings). Interestingly, SCFA treatment markedly inhibited vitamin D-induced osteoclast differentiation **(Figures 12C-E).** Together, these results suggest that vitamin D supplementation promotes a favourable gut microbial composition enriched in SCFA-producing bacteria, and the elevated SCFAs, in turn, suppress vitamin D-mediated osteoclastogenesis, thereby counterbalancing its pro-osteoclastogenic effects.

**Fig. 12:**
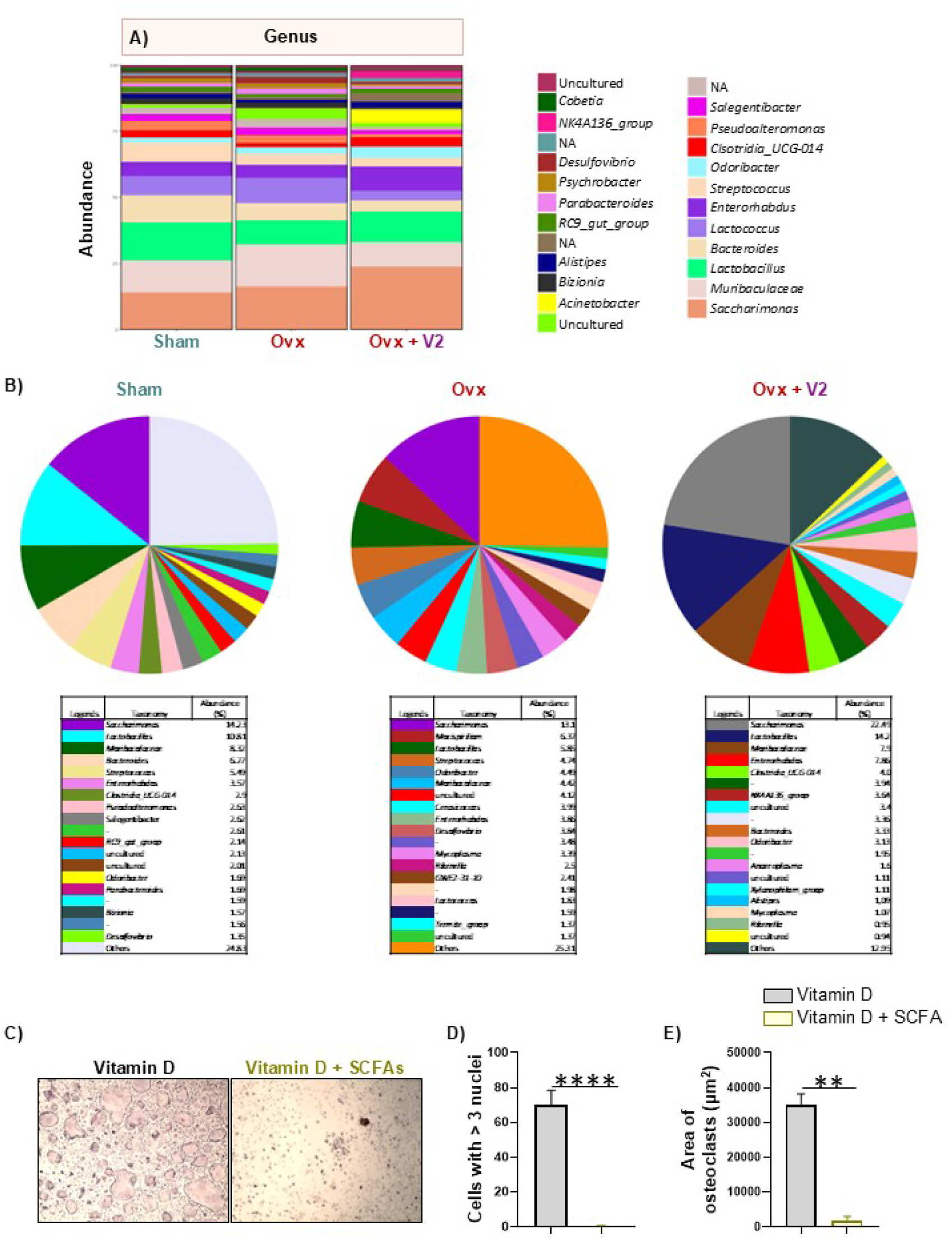
Vitamin D administration enhances abundance of Lactobacillus genus in ovx mice: **(A)** Stacked bar chart representing the relative abundance of the genus in each group. **(B)** Pie charts representing the relative abundance of the genus in each group. BMCs were harvested from mice and were cultured with vitamin D (50 nM) in the presence and absence of short-chain fatty acids (SCFAs) mixture consisting of 0.5 mM of acetate, propionate, butyrate and valerate, along with MCSF (30 ng/ml) and RANKL (60 ng/ml). **(C)** Pictograph images (10X) representing osteoclastogenesis in the presence and absence of SCFAs. **(D)** Bar graph representing the number of multinucleated cells with more than three nuclei. **(E)** Bar graph representing the area of a multinucleated osteoclast. Data are represented as Mean ± SEM. The results were evaluated by Student’s t-test for paired or non-paired data. Statistical significance was defined as *p < 0.05, **p < 0.01, ***p < 0.001 and ****p < 0.0001, concerning the indicated mouse group.

## 4.0 Discussion

Vitamin D plays a crucial role in maintaining bone health; however, its precise mechanisms in regulating bone homeostasis remain incompletely understood due to contradictory reports in the literature. Analysis of vitamin D levels in human PMO subjects revealed a positive correlation with BMD and a negative correlation with both T-score and serum CTX levels, indicating a crucial role of vitamin D in maintaining bone health. To further elucidate the role of vitamin D in preventing osteoporosis, we employed a preclinical ovx mouse model for PMO, and administered vitamin D at both low and high concentrations to study the dose-dependent effect of vitamin D. Our results demonstrated that vitamin D treatment significantly prevented bone loss in ovx mice and enhanced bone strength. To uncover the mechanisms underlying these protective effects, we conducted a series of immunological and molecular analyses.

Since bone loss in osteoporosis largely results from increased osteoclastogenesis, we first examined the direct influence of vitamin D on osteoclast differentiation. *In vitro* assays revealed that vitamin D significantly promoted osteoclastogenesis, consistent with earlier findings that it can enhance bone resorption. This posed an intriguing question: if vitamin D stimulates osteoclast formation *in vitro*, how does it prevent bone loss *in vivo*? To address this, we explored additional alternative pathways through which vitamin D could exert bone-protective effects.

Serum cytokine analysis indicated that vitamin D reduces pro-inflammatory cytokines (IL-6, TNF-α and IL-17) elevated in ovx mice while upregulating anti-inflammatory cytokines (IL-10 and TGF-β), suggesting an immunomodulatory mechanism. Our previous work has established that the immune system critically contributes to osteoporosis pathogenesis, a concept we have defined as “immunoporosis”^23–26^. Building on this foundation, we hypothesized that vitamin D prevents bone loss primarily by mitigating inflammation-induced bone resorption, emphasising the interplay between osteogenic and immune pathways in mediating its osteoprotective effects.

To test this, we investigated the effects of vitamin D on both innate and adaptive immune cells. Within the innate immune compartment, we focused on ILC3s, which are known to express the VDR and regulate cytokines such as IL-17 and IL-22 secretion from ILC3s. We observed that vitamin D did not significantly alter ILC3 frequency but, at higher concentrations, enhanced IL-22 secretion. Furthermore, *in-vitro* analyses showed that vitamin D-primed ILC3s promote osteoclastogenesis in an IL-22-dependent manner.

Next, we examined the adaptive immune cells. Vitamin D administration had negligible effects on Th1 and Th2 subsets but reduced Th17 frequency and increased IL-22 secretion from Th17 cells at higher doses. Interestingly, while Th17 cells are generally pro-osteoclastogenic, vitamin D treatment further enhances their osteoclastogenic potential. Conversely, vitamin D significantly expanded Treg cell populations, with enhanced IL-10 secretion, particularly at higher concentrations. Co-culture experiments confirmed that vitamin D-induced Tregs potently suppressed osteoclastogenesis, implicating Tregs as a key mediator of vitamin D’s bone-protective effects.

We further evaluated the impact of vitamin D on Bregs, which our group has previously reported to inhibit bone loss by suppressing osteoclastogenesis and modulating Treg-Th17 balance^20,22^. Vitamin D administration markedly increased the frequency and anti-osteoclastogenic activity of Bregs at both concentrations, indicating that Bregs may represent an upstream regulatory node through which vitamin D confers bone protection. Further analysis of human subjects revealed that serum vitamin D levels correlate positively with both Breg and Treg cell populations and negatively with Th17 cells, further underscoring the immunomodulatory potential of vitamin D.

Additionally, because IL-22 is known to maintain gut barrier integrity and gut permeability alterations are linked to osteoporosis, we next investigated the role of vitamin D in maintaining the “gut-immune-bone” axis. Vitamin D treatment significantly improved gut barrier function, reducing intestinal permeability in ovx mice, especially at higher doses. Colon and MLNs revealed that vitamin D increased IL-22 production primarily from ILC3s, suggesting that vitamin D’s effects on gut barrier maintenance are mediated through ILC3-derived IL-22. Microbiome profiling further demonstrated that vitamin D supplementation mitigates ovx-induced dysbiosis by enriching beneficial genera such as *Lactobacillus* and *Clostridium*, which are well-established producers of SCFAs^27,28^. Notably, although vitamin D treatment led to an overall reduction in α-diversity, this likely reflects a selective enrichment of functionally beneficial microbial populations rather than a detrimental loss of diversity. These findings suggest that vitamin D reshapes the gut microbiota toward a more favorable and metabolically active composition rather than simply increasing diversity. Interestingly, SCFAs were further observed to counteract the pro-osteoclastogenic effects of vitamin D, indicating a potential feedback interaction between vitamin D signaling, gut microbiota, and bone remodeling.

In summary, our study demonstrates that vitamin D prevents bone loss through multiple interrelated mechanisms: modulation of immune cell balance (enhancing Tregs and Bregs), preservation of gut barrier integrity via IL-22, and restoration of a healthy gut microbiota composition. Collectively, these findings reveal that the overall bone-protective effect of vitamin D results from the convergence of distinct immunological and microbiome-mediated pathways. However, our study has certain limitations. Among the innate immune cells, we primarily focused on ILCs and did not investigate the roles of macrophages, dendritic cells, or neutrophils. The influence of vitamin D on these immune cell populations could also contribute to the regulation of osteoclastogenesis and, consequently, bone homeostasis. Our analyses have overlooked vitamin D’s regulatory effects on osteoblastogenesis, a critical pathway for bone formation. Furthermore, immune cell analysis was performed on a relatively small cohort of patients, and although significant associations between serum vitamin D levels and bone health parameters were observed, these findings remain correlative and do not establish causality. While efforts were made to minimize bias by excluding participants with major metabolic disorders or those receiving medications known to affect bone metabolism or vitamin D status, other potential confounding factors, such as dietary intake, sunlight exposure, and lifestyle variables, were not fully controlled or adjusted for in the statistical analysis. Additionally, future studies employing VDR knockout mice would provide deeper insights into the molecular mechanisms underlying vitamin D-mediated regulation of bone health.

Conclusively, our findings indicate that vitamin D preserves bone in estrogen deficiency not merely by supporting mineral metabolism, but through coordinated immunomodulation and maintenance of the gut-immune-bone axis. This integrated mechanism reconciles the apparent paradox of pro-osteoclastogenic effects *in vitro* with net anti-resorptive outcomes *in vivo* and nominates immune-gut microbial pathways as leverage points for optimizing vitamin D-based strategies in osteoporosis prevention. Our findings from the present study open novel therapeutic avenues that via restoring the gut-microbial composition, vitamin D mitigates inflammatory bone loss even under estrogen-deficient PMO conditions.

## Acknowledgment

AB, LS, SR, AS, SY, CS and RKS acknowledge the Department of Biotechnology and Central Core Research Facility (CCRF), AIIMS, New Delhi, India, for providing infrastructural facilities. AB and SR thank ICMR for the research fellowship. SY thanks the DBT for the research fellowship. CS thanks the CCRH-Ayush for the Research Fellowship.

## Conflict of Interest Statement

The authors declare no conflicts of interest.

## Funding

This work was financially supported by projects: Indian Council of Medical Research ICMR (61/05/2022-IMM/BMS) and ICMR (EMDR/IG/13/2024-01-00842) sanctioned to RKS.

## Credit authorship contribution statement

RKS contributed to the conception and design of the study; AB, LS and TS contributed to the acquisition and analysis of data; AB contributed to drafting the text or preparing the figures. SR, PS and AWB helped with H & E and TRAP staining of bone tissues. SY and AS helped with micro-CT. CS helped with RT-PCR. BG and VM provided human samples. PKM provided valuable inputs. All authors reviewed the manuscript. All authors contributed to the article and approved the submitted version.

## Data Availability Statement

16S rRNA gene sequencing data generated in this study has been deposited in the Mendeley data repository (https://data.mendeley.com/research-data/?query=10.17632/7j57nk95sv.1).

**<u>Send correspondence to</u>:**

Dr. Rupesh K. Srivastava,

Additional Professor,

*Translational Immunology, Osteoimmunology & Immunoporosis Lab (TIOIL)*

*An ICMR Collaborating Centre of Excellence on Bone Health*

Department of Biotechnology,

All India Institute of Medical Sciences (AIIMS),

New Delhi-110029, India

Cell: +91-9179567399

**Supplementary Fig. 1:**
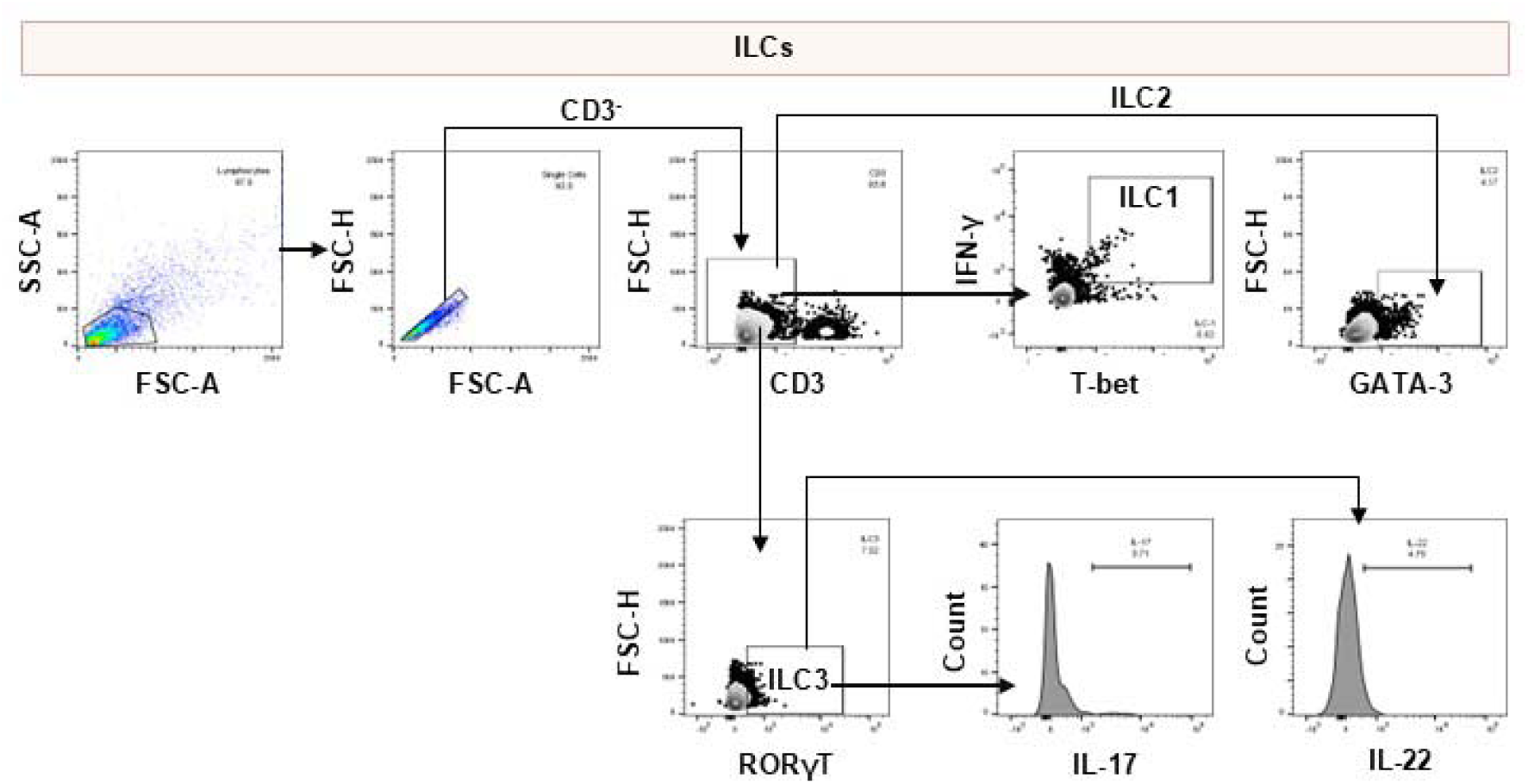
Gating strategy used for the analysis of ILCs.

**Supplementary Fig. 2:**
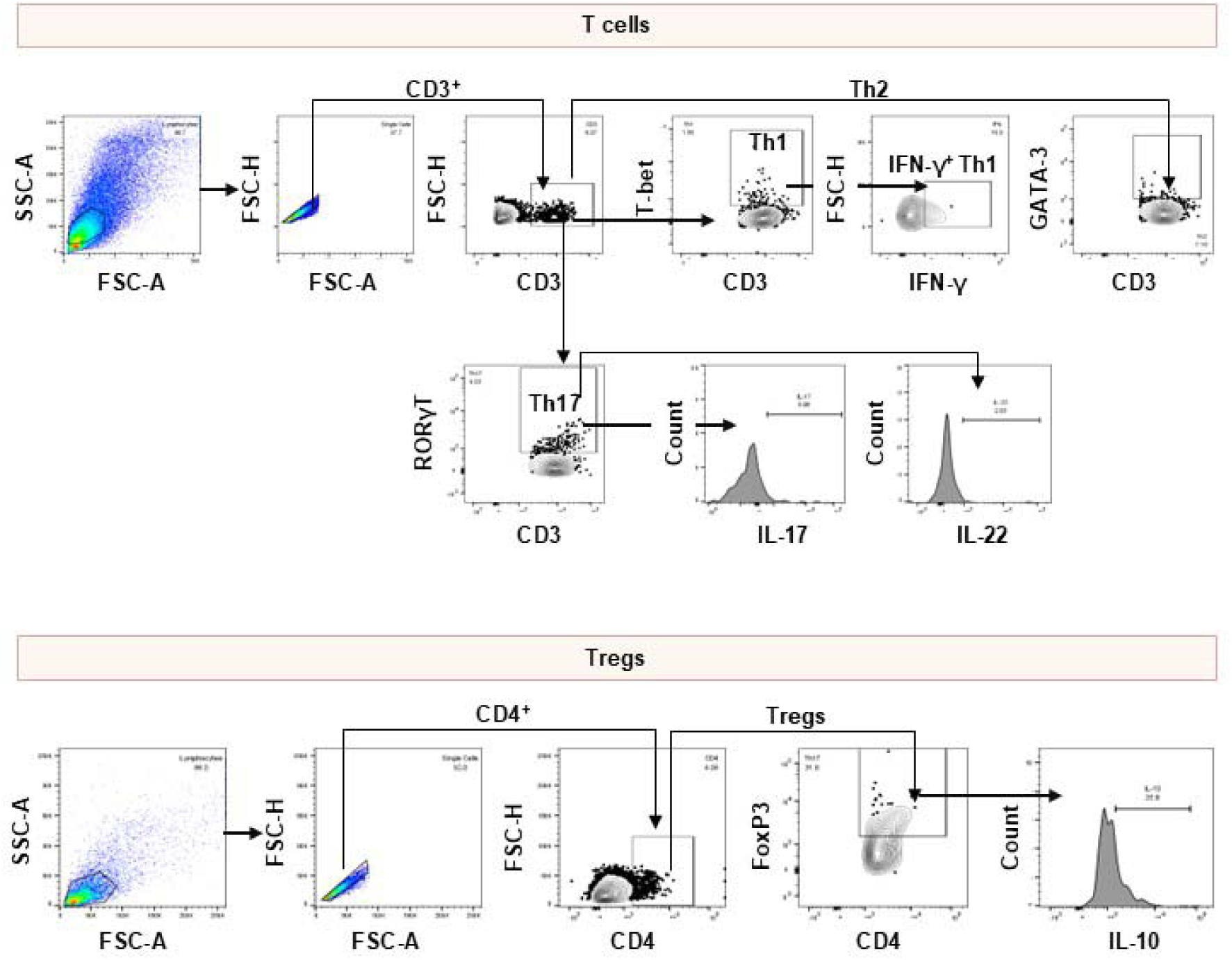
Gating strategy used for the analysis of T cells.

**Supplementary Fig. 3:**
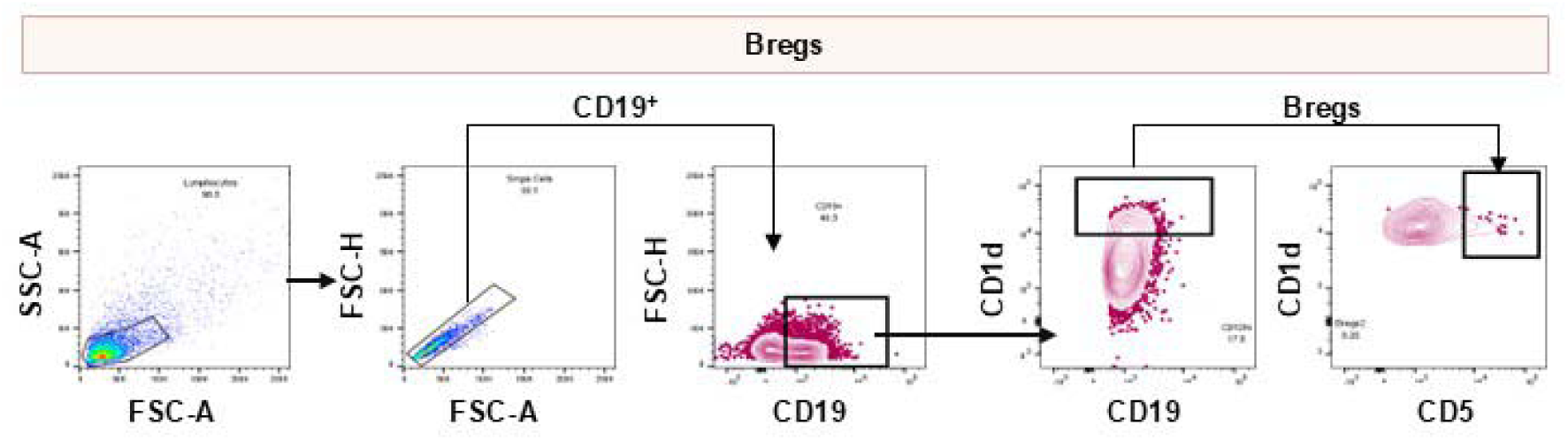
Gating strategy used for the analysis of Bregs.

**Supplementary Fig. 4:**
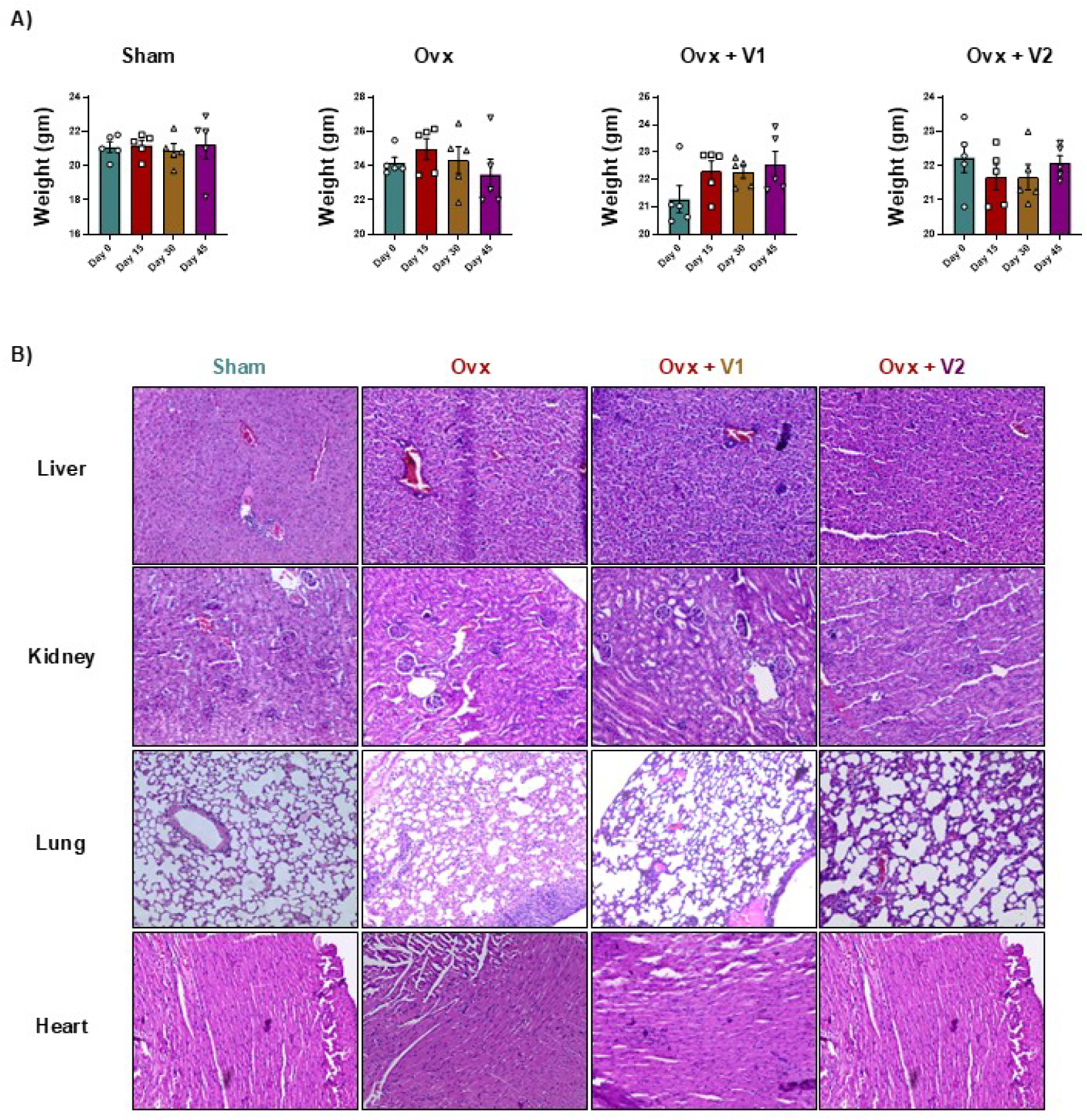
Assessment of systemic toxicity following vitamin D treatment. **(A)** Body weight of mice in sham, ovariectomized (ovx) and vitamin D-treated groups (V1: 2 ng/mouse/day; V2: 4 ng/mouse/day) at different time points. **(B)** Representative histological images (H&E staining) of major non-skeletal organs, including liver, kidney, lung, and heart. Data are represented as Mean ± SEM. The results were evaluated by one-way ANOVA followed by Dunnett’s test. Statistical significance was defined as *p < 0.05, **p < 0.01, ***p < 0.001 and ****p < 0.0001, concerning the indicated mouse group.

**Supplementary Fig. 5:**
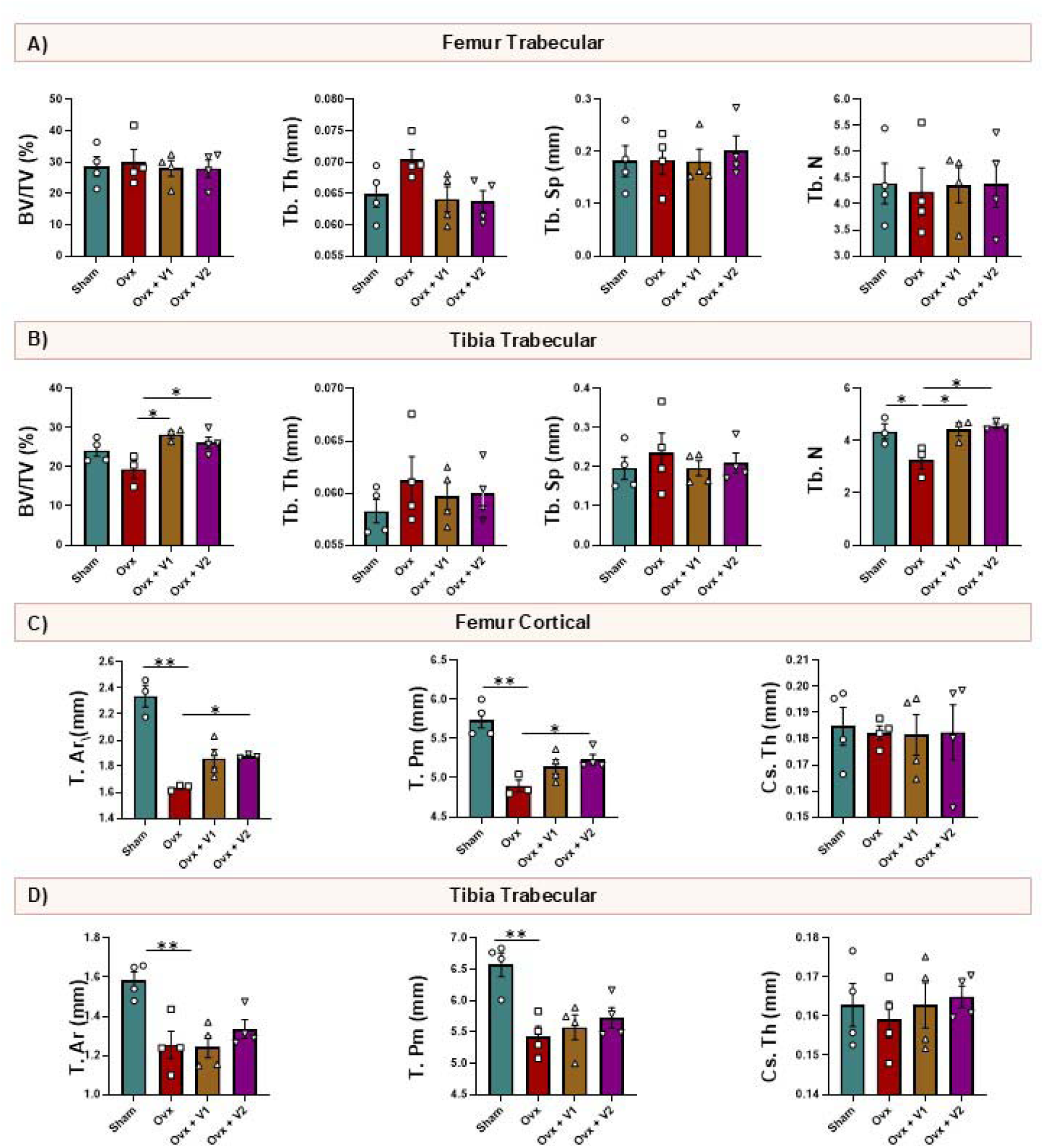
Vitamin D administration prevents bone loss in ovx mice. Histomorphometric parameters of the (**A)** femur trabecular bone, **(B)** tibia trabecular bone, **(C)** femur cortical bone and **(D)** tibia cortical bone. Bone volume/tissue volume ratio; Tb.Th, trabecular thickness; Tb.Sp, trabecular separation, Tb. N, trabecular number; T. Ar, total cross-sectional area; T. Pm, total cross-sectional perimeter; Cs. Th, cross-sectional thickness. Data are represented as Mean ± SEM. The results were evaluated by one-way ANOVA followed by Dunnett’s test. Statistical significance was defined as *p < 0.05, **p < 0.01, ***p < 0.001 and ****p < 0.0001, concerning the indicated mouse group.

**Supplementary Fig. 6:**
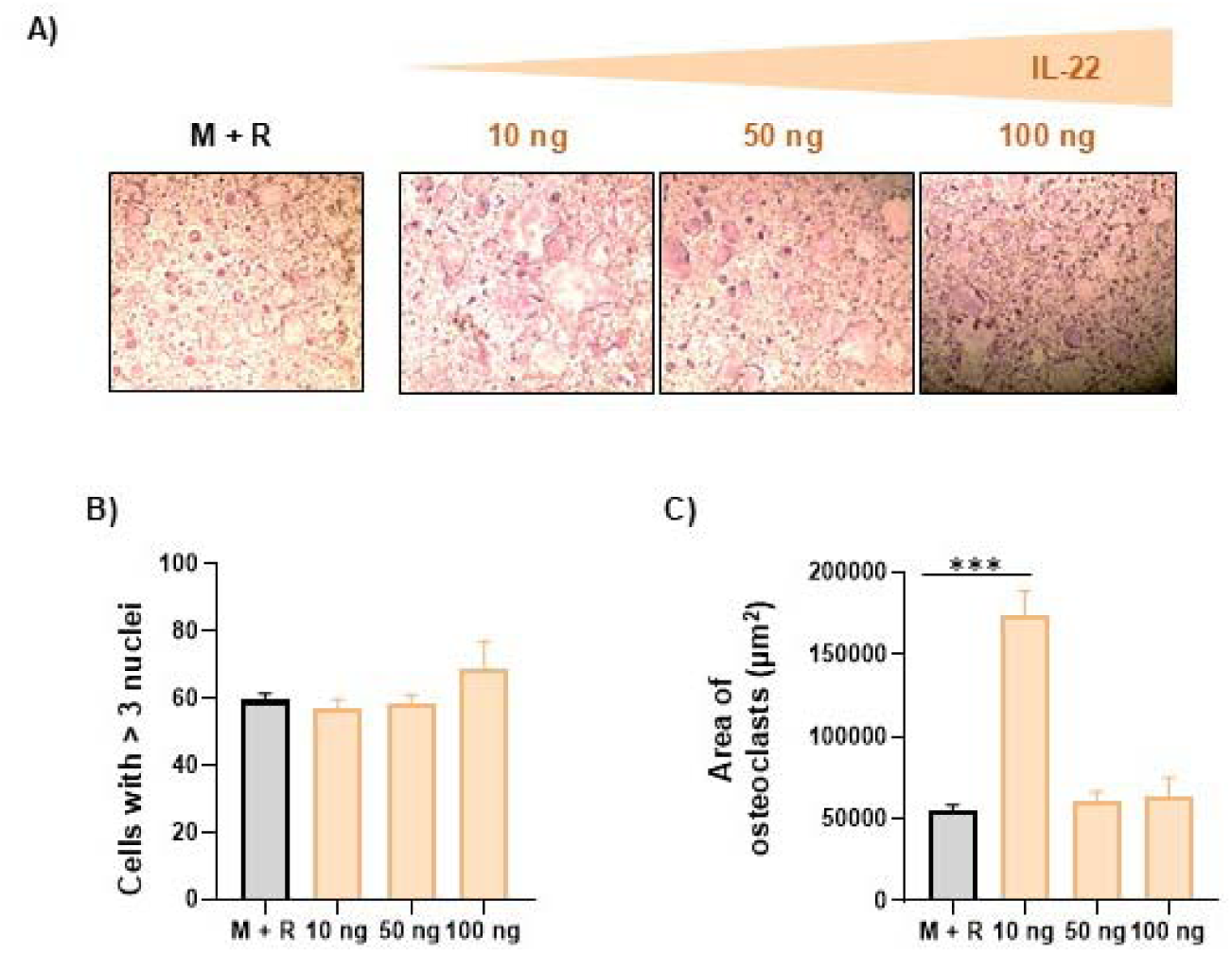
IL-22 promotes osteoclastogenesis of bone marrow cells (BMCs). BMCs were harvested from mice and were cultured in the presence or absence of IL-22 along with MCSF (30 ng/ml) and RANKL (60 ng/ml). **(A)** Pictograph images (10X) representing osteoclastogenesis in the presence or absence of IL-22. **(B)** Bar graph representing the number of multinucleated cells with more than three nuclei. **(C)** Bar graph representing the area of a multinucleated osteoclast. Data are represented as Mean ± SEM. The results were evaluated by one-way ANOVA followed by Dunnett’s test. Statistical significance was defined as *p < 0.05, **p < 0.01, ***p < 0.001 and ****p < 0.0001, concerning the indicated mouse group.

**Supplementary Fig. 7:**
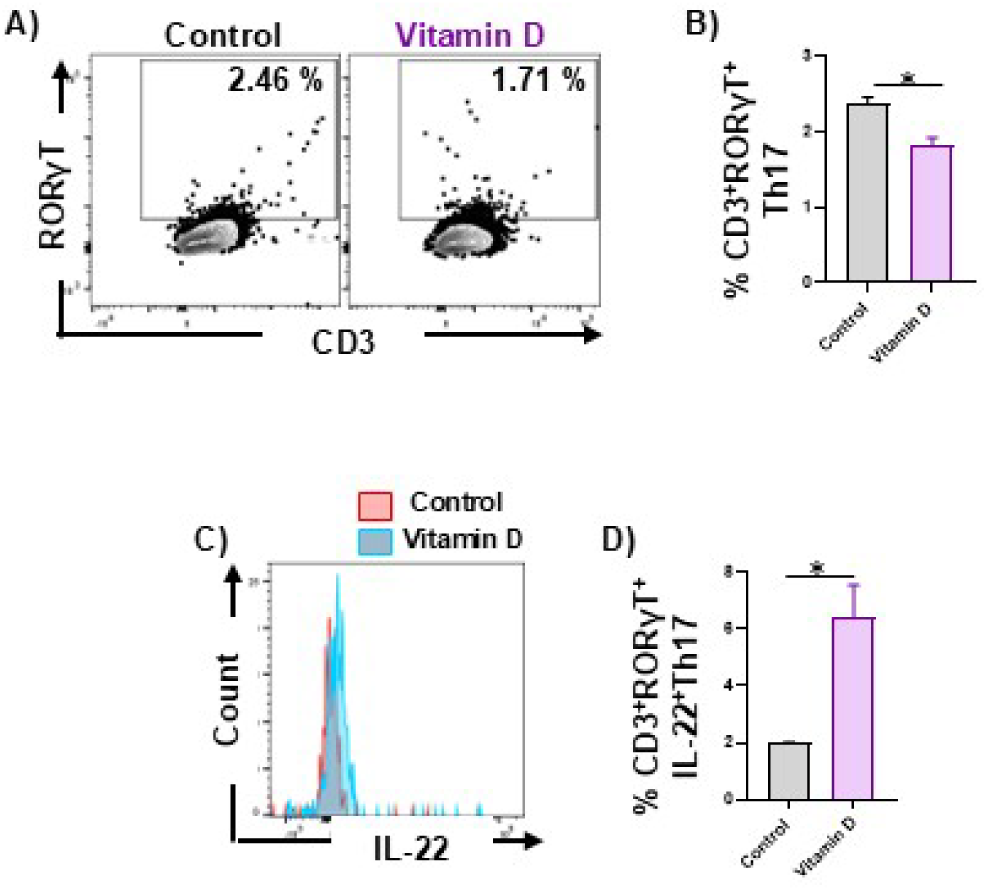
Vitamin D inhibits differentiation of Th17 cells *in vitro*. Naïve T cells were cultured under Th17-polarising conditions in the presence or absence of vitamin D (50 nM) for 5 days. After 5 days, **(A & B)** Th17 and **(C & D)** IL-22^+^ Th17 cells were analyzed in the cultured cells with the help of flow cytometry. Data are represented as Mean ± SEM. The results were evaluated by Student’s t-test for paired or non-paired data. Statistical significance was defined as *p < 0.05, concerning the indicated mouse group.

**Supplementary Fig. 8:**
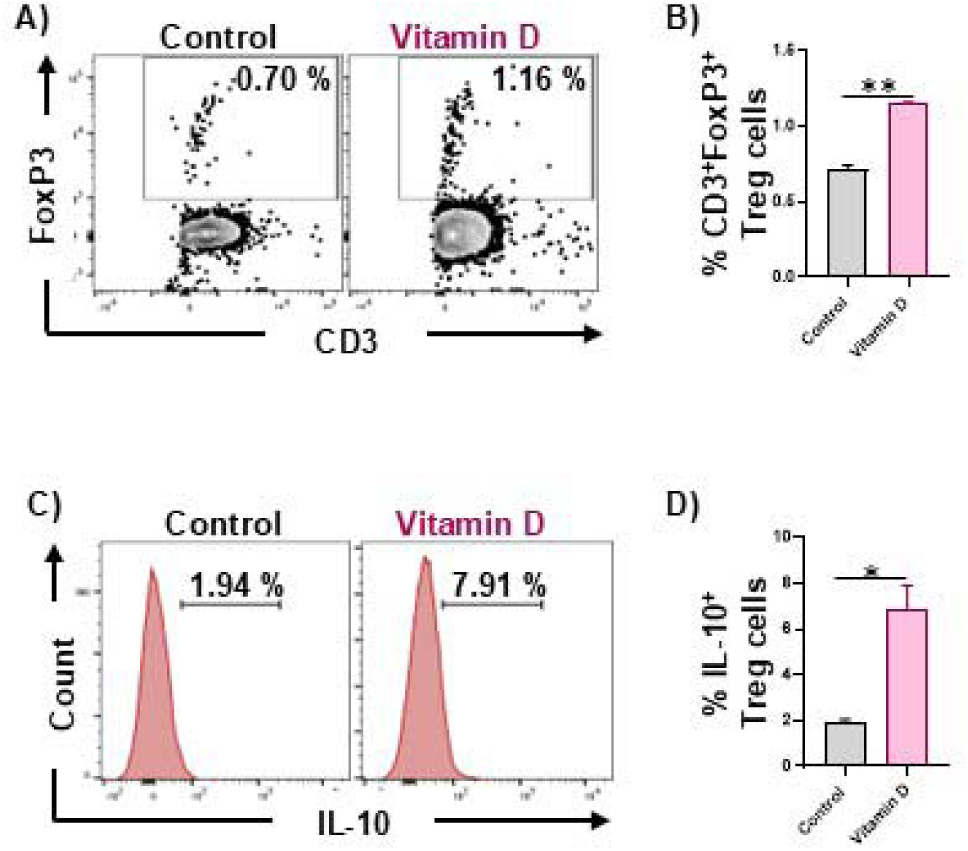
Vitamin D promotes differentiation of Treg cells *in vitro*. Naïve T cells were cultured under Treg-polarising conditions in the presence or absence of vitamin D (50 nM) for 5 days. After 5 days, **(A & B)** Treg and **(C & D)** IL-10^+^ Treg cells were analyzed in the cultured cells with the help of flow cytometry. Data are represented as Mean ± SEM. The results were evaluated by Student’s t-test for paired or non-paired data. Statistical significance was defined as *p < 0.05, and **p < 0.01, concerning the indicated mouse group.

**Supplementary Fig. 9:**
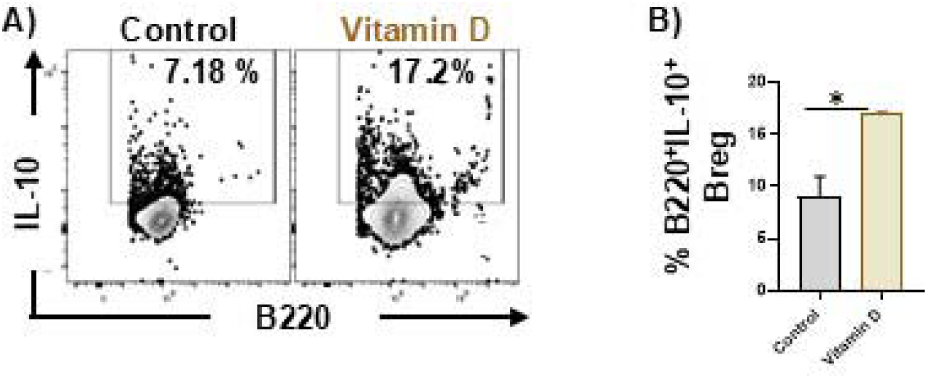
Vitamin D promotes differentiation of Bregs *in vitro*. B cells were cultured in the presence or absence of vitamin D (50 nM) for 24 hrs. **(A & B).** After 24 hrs, Bregs were analyzed in the cultured cells with the help of flow cytometry. Data are represented as Mean ± SEM. The results were evaluated by Student’s t-test for paired or non-paired data. Statistical significance was defined as *p < 0.05, concerning the indicated mouse group.

## Notes

### Competing Interest Statement

The authors have declared no competing interest.

